# Phytochrome light receptors control metabolic flux, and their action during seedling development sets the trajectory for biomass production

**DOI:** 10.1101/762666

**Authors:** Johanna Krahmer, Ammad Abbas, Virginie Mengin, Hirofumi Ishihara, Thiago A Moraes, Nicole Krohn, Maria-Grazia Annunziata, Regina Feil, Saleh Alseekh, Toshihiro Obata, Alisdair R Fernie, Mark Stitt, Karen J. Halliday

## Abstract

The phytochromes (phys) photoreceptors are known to be major regulators of plastic growth responses to vegetation shade. Recent reports have begun to uncover an important role for phys in carbon resource management. Our earlier work showed that *phy* mutants had a distinct metabolic profile with elevated levels of metabolites including TCA intermediates, amino acids and sugars. Here we show that in seedlings phy regulates the balance between glucose and starch. Multi-allele *phy* mutants have excess glucose and low starch levels, which is conducive to hypocotyl elongation. ^13^C-CO_2_ labelling demonstrates that metabolic flux balance in adult plants is markedly altered in *phy* mutants. Phytochrome reduces synthesis rates of stress metabolites, including raffinose and proline and several typical stress-induced biosynthetic genes related to these metabolites show higher expression in phy mutants.

Since growth and metabolism are typically inter-connected, we investigated why *phy* mutants have severely reduced biomass. Quantification of carbon fixation, biomass accumulation, and ^13^C labelling of cell wall polysaccharides established that relative growth rate is impaired in multi allele *phy* mutants for the first 2.5 weeks after germination but equivalent to the WT thereafter. Mathematical modelling predicts that the altered growth dynamics and final biomass deficit can be explained by the smaller cotyledon size of the multiple *phy* mutants. This indicates that the established role of phy in promoting seedling establishment has enduring effects that govern adult plant biomass.

## Introduction

Phytochrome light receptors are major regulators of growth plasticity, a fundamental characteristic that ensures plants adapt to a changing environment. The shade avoidance response (SAR) is a common adaptive growth strategy in vegetation-rich habitats, where competition for light and other resources can be intense. In Arabidopsis, a rosette plant, SAR features include increased petiole elongation and a reduced leaf blade area. These large changes in leaf architecture require modifications of leaf development and carbon resource management (Yang et al. 2016). While we have a good appreciation of the molecular events that underlie the SAR, our knowledge of the concomitant metabolic changes is more rudimentary.

In nature, the light that a plant experiences through a day depends on its proximity to objects that occlude light and in particular its proximity to other plants. Dense vegetation can restrict access to light through shading, while the absorbance properties of plant pigments also elicit changes in light spectral quality resulting in a higher proportion of far-red (FR) compared to red (R) wavelengths. The phytochrome photoreceptors are exquisitely tuned to detect these vegetation-induced changes in light quantity and quality. Phytochromes exist in two isoform states, Pr (R-absorbing) and Pfr (FR-absorbing), which allows them to operate as biological light switches. R light induces Pr photoconversion to the active Pfr form, while FR switches Pfr back to inactive Pr. Thus, changes in the R:FR ratio within vegetation-rich habitats drive the dynamic equilibrium of Pr:Pfr, and the proportion of active Pfr (Rausenberger et al. 2010). A property known as dark or thermal reversion, which leads to light independent relaxation of Pfr to Pr, is important for phytochrome detection of light fluence rate, particularly under shaded conditions. The low R:FR, and potentially low irradiance conditions of dense vegetation inactivate phytochrome and elicit an adaptive SAR growth strategy (Rausenberger et al. 2010).

In seedlings, SAR typically results in elongation of hypocotyls, delayed apical hook opening, reduced cotyledon size, and low chlorophyll content (Hu et al. 2013; Chen and Chory 2011; K. A. Franklin and Whitelam 2005; Leivar et al. 2008). In adult plants SAR features include elongated petioles, smaller leaf blades, increased leaf hyponasty, reduced leaf emergence rate and early flowering (Casal 2013; Halliday 2003). In Arabidopsis, there are five phytochromes designated phyA-phyE. PhyB is known to have a central role in regulating SAR, indeed *phyB* mutants have a constitutive SAR phenotype (Reed et al. 2007). Higher order mutants that lack other phytochromes in addition to phyB, e.g. *phyBD, phyABD* and *phyABDE*, exhibit incrementally more severe SAR phenotypes (Yang et al. 2016).

We have a growing understanding of the molecular and hormonal signalling pathways that underlie SAR. Active Pfr suppresses the activity and / or expression of the PIF (PHYTOCHROME INTERACTING FACTOR) transcription factors, therefore PIFs are derepressed in the shade (Leivar et al. 2012). In seedlings, PIF4, PIF5 and PIF7 mediate SAR under a low R:FR ratio, low irradiance light, through sequential auxin biosynthesis and signalling phases. Detailed analysis showed that mesophyll cell PIF4 levels correlate with the early rise and subsequent fall of auxin levels. In prolonged low R:FR ratio, PIF4 levels remain elevated in epidermal and sub-epidermal hypocotyl cells, which appears to enhance auxin perception (via ABF3, TIR1 and MIR393) and auxin signalling (via *IAA19, IAA29*). In adult plants PIF7 appears have a dominant role in regulating SAR, again through auxin biosynthesis and signalling (Mizuno et al. 2015; de Wit, Ljung, and Fankhauser 2015).

What is becoming increasingly clear is that phytochrome action is interwoven with carbon metabolism and signalling. In Arabidopsis seedlings, phyB and PIF1, PIF3, PIF4 and PIF5 have been shown to have regulatory functions in sucrose promotion of hypocotyl elongation (Stewart, Maloof, and Nemhauser 2011; Lilley-Steward et al. 2012). Phytochromes are proposed to be involved in promoting root growth by cotyledon-derived sucrose (Kircher and Schopfer 2012). Work in *Brassica* seedlings illustrates that the proportion of assimilated carbon that is partitioned to the hypocotyl doubles in response to low R:FR conditions (de Wit et al. 2018). This low R:FR induced change in carbon allocation is mainly dependent on PIF7, with contributions from PIF4 and / or PIF5. Phloem transport of sucrose by the SUC2 and SWEET11 and SWEET 12 transporters is necessary to fuel hypocotyl elongation in response to low R:FR (de Wit et al. 2018). Thus in seedlings phys are not only required for altered molecular signalling but also for reprogramming of carbon resource management.

Our published work, and that of others has shown that in *phy* mutants a broad spectrum of metabolites have altered levels (Yang et al. 2016). Sugars, TCA cycle components and amino acids, in particular, are more abundant in multi-allele *phy* mutants (Yang et al. 2016). We also established that in addition to the well described changes in leaf morphology, phy deficiency compromises biomass production, leading to an 80% reduction in *phyABDE* mutant rosettes (Yang et al. 2016). However the regulatory processes that underly phy-control of metabolite levels and plant biomass are currently unknown.

In this paper we analyse the impact of phy-deficiency on metabolic flux using 13C labelling, LC-MS and GC-MS metabolite profiling methods. Our data show that phytochromes play a pivotal role in carbon-resource management, with distinct functions in early and later development. In seedlings phytochromes regulate the balance between glucose and starch, which is important for hypocotyl elongation. In adult plants phytochromes control *de novo* synthesis of stress metabolites such as proline and raffinose. By using the multi-scale Framework Model, that predicts environmental control of leaf and whole plant growth, we have developed a new system-level understanding of why phy-depletion has such a dramatic impact on plant biomass.

## Results

### phy control of metabolism is developmental stage-dependent

We have previously observed substantial increases in sugars, organic acids and amino acids in adult phytochrome mutants (Yang et al. 2016). In particular, malate, proline starch and glucose were consistently high in severely phy-deficient mutants. We rationalized that this may not be the case in phy-depleted seedlings that have much lower chlorophyll levels (Hu et al. 2013) and therefore potentially lower rates of carbon assimilation. End of day (EoD) measurements with 10 and 14 day-old seedlings established that multi-allele mutants *phyABD*, and particularly *phyABDE*, had lower levels of malate, proline and starch than WT (Ler) seedlings (Figure 1A-C). However, glucose over-accumulated in the *phy* mutant seedlings, like in older rosettes (Figure 1D) (Yang et al. 2016). Since we observed that seedlings but not older rosettes *phy* mutants have a high water content (Figure S1A, B) we re-normalized metabolite abundance on DW. While this attenuated the differences between WT and *phy* mutants, the results were qualitatively similar to the fresh weight normalisation (Figure S1C-F; Figure 1).

**Figure 1:**
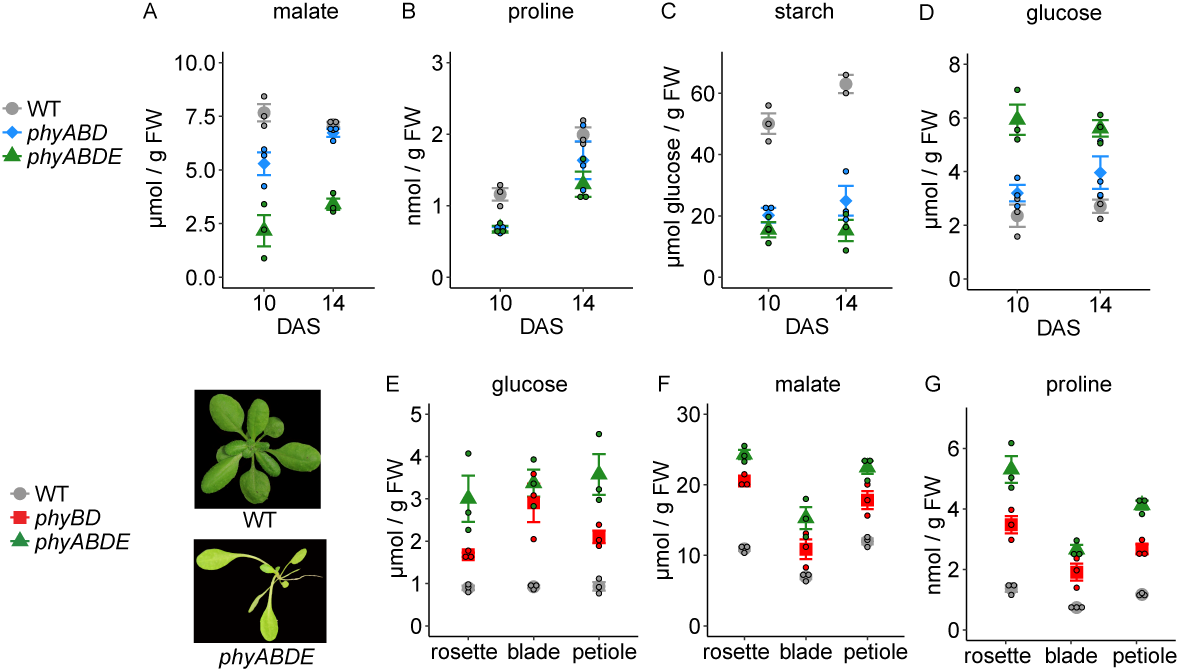
Metabolite content of phy mutants vs WT as seedlings and different tissues of adult plants. (A-D) Selected metabolites measured in WT (Ler), *phyABD* and *phyABDE* seedlings at 10 DAS and 14 DAS. A) malate, B) proline C) starch D) glucose. (E-G) Metabolite content of adult WT, phyBD and phyABDE, measuring from either entire rosettes, leaf blades, and petiole enriched tissue. (E) glucose, (F) malate, (G) proline. Conditions were LD 12:12, 115µE, 18°C in all experiments. Legend symbols indicate mean, errorbars SEM, coloured circles represent the individual measurements of replicates.

Given this developmental-stage conditionality, we wanted to establish whether the metabolite excess was a consistent feature of adult phy-deficient mutants. Plants were subjected to a range of growth regimes including varying photoperiods (8L:16D, 12L:12D and 16L:8D), irradiance levels (54, 110 or 190 μE) or temperatures (16°C or 22°C) in 12L:12D (Figure S2A-I). Light and temperature changes were applied only for the last 3 days of the experiment, after growth in standard conditions (18°C, 115µE) to ensure that plants of a similar size and developmental stage are compared. Metabolites were sampled at EoD. In all conditions we observed higher levels of glucose, malate and proline in *phy* multi-allele mutants compared to WT, the only exception being *phyBD* in 8L:16D which had similar glucose levels to WT (Figure S2A-I). Enhanced levels of glucose, malate and proline were also observed in the Ler and Columbia *phyABD* mutant lines compared to the respective wild-types, indicating that the effect was not accession-specific.

Since a large proportion of responses to low phy activation are mediated by increased activity of the transcription factors PIF4 and PIF5, we measured a selection of metabolites in p35S:PIF4-HA and p35S:PIF5-HA (Lorrain et al. 2008). Both lines showed increased metabolite levels, but this was not the case for the *hy5* mutant, a positive phy signalling element (Fig S2M-O).

As phy-deficient mutants have altered leaf morphology, with longer petioles and smaller blades, we checked whether metabolism might differ between these two tissues. We found that in rosette, leaf blade and petiole tissue, metabolite levels were consistently highest in the quadruple *phyABDE* mutant, followed by *phyBD*, then WT (Figure 1E-G). Collectively, our data show that in adult plants the impact of phytochrome depletion on glucose, malate and proline levels is extremely robust to environmental perturbation and is widespread across vegetative tissues. Also, for some metabolites there is a switch from under to over accumulation through *phy* mutant development.

### LC-MS/MS analysis reveals phy-dependent changes in phosphorylated sugars and other central metabolites and the sucrose-signal T6P

To gain a deeper understanding of how phytochrome influences primary metabolism, we used LC-MS/MS, as it provides a reliable method to quantify phosphorylated components and other metabolites of central metabolism (Arrivault et al. 2009; Szecowka et al. 2013). As nany of these metabolites have extremely short half-lives, plant material was rapidly quenched under growth irradiances at 6h into the photoperiod and without exposure to light at the end of the night (24h) in mature 35 day old 12L:12D-grown *phyABD* and WT plants (Figure 2a,c). We included samples from 30 day-old WT plants, with the same biomass as the *phyABD* to control for the possibility that plant size influences metabolite abundance.

**Figure 2:**
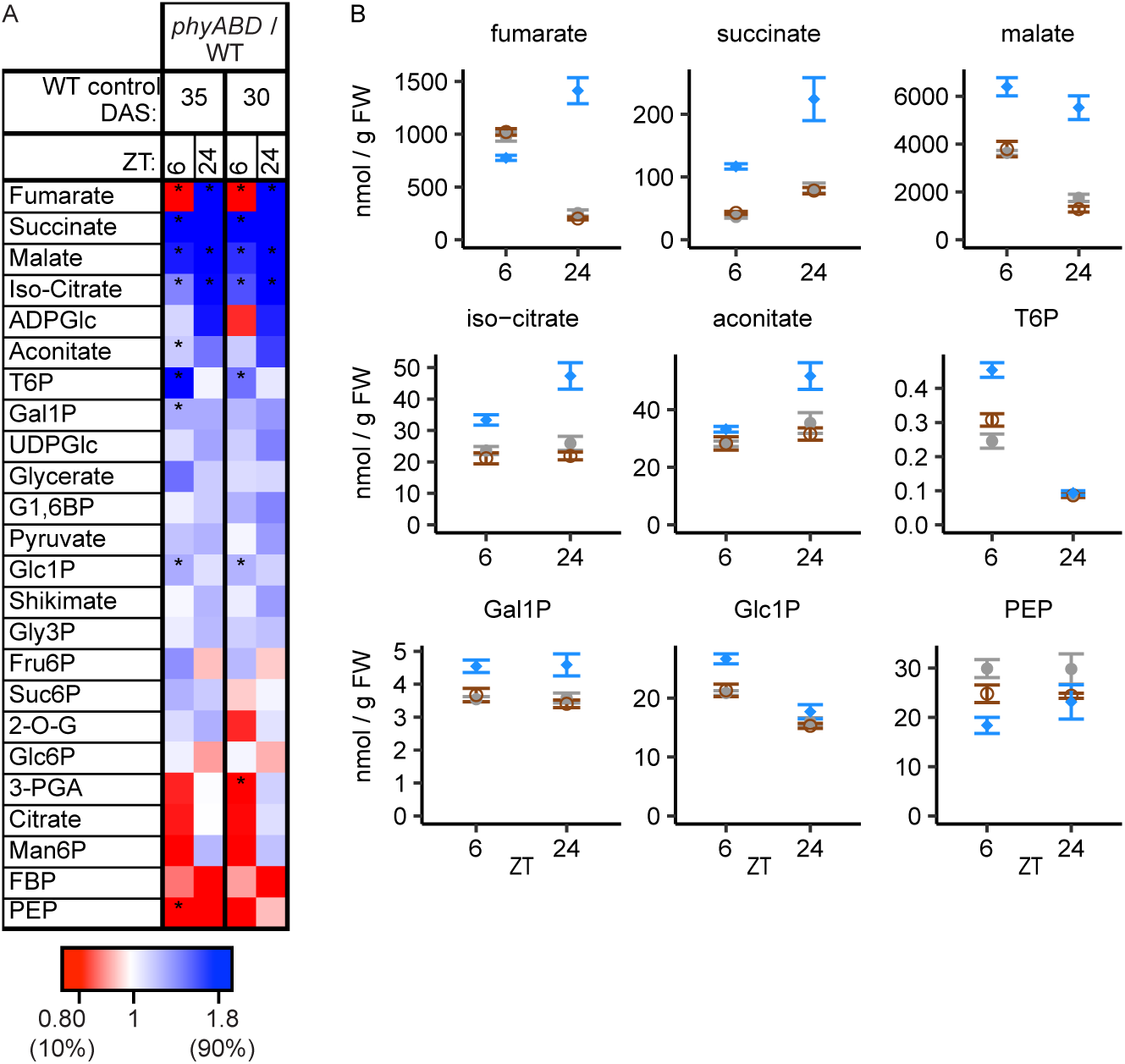
Metabolite abundance measured by LC-MS at ZT6 and ZT24 in *phyABD* at 35 DAS, a same age WT control (35 DAS) and a same biomass WT control (30 DAS). A) Heatmap representing the fold change in *phyABD* over either of the WT controls. Blue: fold change > 1, red: fold change < 1. * p<0.05 in t-test comparing WT and *phyABD* of the same time point. B) Plots of abundance of metabolites with a significantly decrease or increased abundance in *phyABD*. Samples were harvested and frozen inside the incubator to preserve phosphorylated and short-lived metabolites.

In line with our published data, in *phyABD* we observed elevated levels of TCA components, succinate, iso-citrate, malate and fumarate (Yang et al. 2016) (figure 2C, Figure S3). Levels of 3-carbon metabolites were similar in *phyABD* and WT, except for PEP and 3-PGA, which were slightly lower in *phyABD* during the daytime. Expected diel changes were observed for many of the phosphorylated sugars, whose levels were higher during the day than the night. Interestingly, daytime levels of T6P, Suc6P and Glc1P were higher in *phyABD*, while Gal1P levels were slightly raised at 6 and 24h time points (figure 2, figure S3). These data indicate that phy depletion leads to a shift in the balance of TCA components and major metabolic sugars, as well as the sucrose-signal T6P (John Edward Lunn et al. 2014; Figueroa and Lunn 2016).

### Phytochrome depletion alters the rate of label incorporation into metabolites

Next we combined time-resolved GC-MS assays with ^13^C-CO_2_ labelling to allow us to track both the diurnal abundance and the flux to individual metabolites. For direct comparison to LC-MS, we again used *phyABD* and WT 35/30 day-old controls, and younger (17-19 d) WT, *phyABD, phyABDE* plants. Metabolites were quantified by GC-MS at intervals through a 12L:12D cycle, and ^13^CO_2_ labelling was performed from ZT0 until ZT2 or ZT12 allowed the measurement of metabolic flux to label saturation (Arrivault et al. 2009; Szecowka et al. 2013) (Figure 3A-B). We estimated label incorporation into individual metabolites by identifying the time point before enrichment saturated, multiplying the enrichment at this time by the abundance of the metabolite at this time., and then normalising the value on that in wild-type plants. This value is an approximate proxy for the minimum rate of synthesis of the metabolite; it will be an underestimate of the rate of synthesis if enrichment in the precursor is less than 100% and if the metabolite is further metabolised.

**Figure 3:**
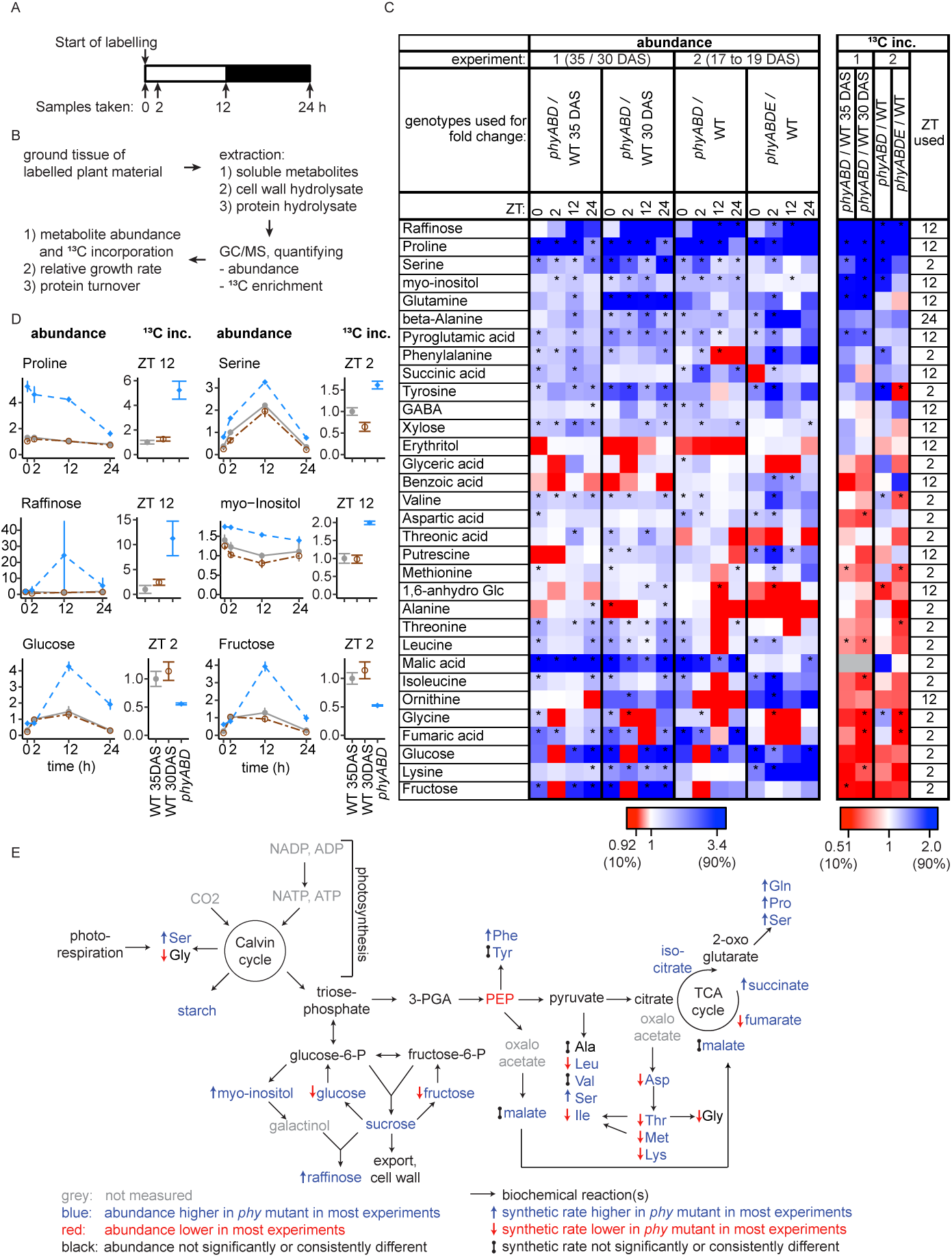
GC-MS analysis of metabolite abundance and ^13^C label incorporation in *phy* mutants and WT controls. A) Timing of ^13^C labeling and sampling. B) Workflow of sample preparation for GC/MS and data analysis. Rate of ^13^C label incorporation was calculated by multiplying the abundance at a given time point by the fraction of ^13^C in the metabolite at the same time point. C) Heatmap displaying fold change in abundance (left) and rate of ^13^C label incorporation (right, ‘^13^C inc.’) of *phy* mutant / WT control. For experiment 1, fold change in *phyABD* is shown compared to the same age and same biomass controls, for experiment 2, fold change over WT is shown for *phyABD* and *phyABDE*. All abundance time points are shown, but only the most suitable time point for the rate of label incorporation (time indicated in the rightmost column), which are the latest time points before label saturation (typically 2 or 12h). Rows are sorted by decreasing average fold change in label incorporation. Blue and red denote higher and lower values in the *phy* ko, respectively. Grey: values that could not be determined. * p < 0.05, two-sided t-test. D) Plots of abundance and label incorporation of selected metabolites from experiment 1. E) Simplified and generalized daytime metabolic pathway map illustrating the observed changes in abundance and ^13^C incorporation.

The heat map (Figure 3C, Figure S4) shows metabolite abundance and ^13^C incorporation as the fold-change ratio a given *phy* mutant and WT. Except for some minor differences, the changes in the *phy* mutants are broadly similar in different aged plants. Consistent with our published data, Fig. 2 and Supplemental Figure 3, *phy* mutants exhibit elevated levels of several organic acids, amino acids, sugars and particularly high levels of malate, and the stress associated metabolites proline and raffinose (Figure 3C-D, Figure S4-7).

The ^13^C labelling data illustrate that the increased levels of the amino acids proline, glutamine and serine in *phy* mutants, could result from their increased rates of synthesis from newly fixed C (Figure 3C, 3). These three amino acids can be synthesized from 2-oxo-glutarate (2-OG), indicating a potential common up-regulation of biosynthetic processes using 2-OG (Figure 3E). As 2-OG is not very amenable to GC-MS analysis, we do not have data on its enrichment. However, its abundance was unaltered in the *phy* mutants according to LC-MS/MS analysis at 6h (figure 2A), in spite of up-regulation of most other TCA-cycle components, which is consistent with use of 2-OG for increased amino acid synthesis. Phy-depletion also increases levels and synthetic rates of raffinose and its precursor myo-inositol (Figure 3C-D), as well as beta-alanine phenylalanine, and glutamine in older plants.

For several metabolites, levels are mildly elevated in *phy* mutants, but this is not obviously due to faster synthesis from newly fixed C. Examples include amino acids in the oxaloacetate and pyruvate branches such as lysine and threonine, where phy-depletion leads to slightly raised levels, but lower or unchanged label incorporation in particular in the old plants (Figure 3C-E). Pronounced abundance changes are observed for malate, glucose and fructose in *phy* mutants compared to WT (Figure 3C-D, S4-7). In the WT, net synthesis of these metabolites is rapid, reaching steady state levels by ZT2. However, the dynamics are quite different in severe *phy* mutants, that have a higher dawn content, little further accumulation in the first 2 h of the light period and ower initial labelling rates but rising levels later in the daytime with two-fold higher levels at dusk than WT plants. This suggests that daytime demand for these metabolites is lower in *phy* mutants, or that they accumulate in the vacuole or other organelles.

In summary, the ^13^C labelling data indicate that phytochrome depletion alters the flux balance to different sets of metabolites with increased label incorporation into amino acids like proline, serine and glutamine, which are derived from 2-OG. Likewise, phy-depletion boosts raffinose and myo-inositol synthesis and abundance, while inducing over-accumulation of glucose and fructose, possibly due to decreased use of these metabolites.

### Phytochrome control of stress metabolites likely involves ABA and *CBF3*

The metabolic profile of *phy* mutants is reminiscent of metabolic changes observed in response to drought, cold stress and ABA treatment (Cook et al. 2004; Urano et al. 2009; Kempa et al. 2008; Pagter et al. 2017). In particular, the two metabolites with the highest over-accumulation and synthetic rate, raffinose and proline, have been described as typical stress-induced metabolites in response to cold treatment or drought in Arabidopsis and other plants e.g. (e.g. Xin & Browse, 1998; Hare & Cress, 1997; Gilmour *et al*, 2000). Therefore we hypothesized that phy depletion may lead to induction of signalling pathways shared with such stress conditions.

Previously we showed that both *phyABDE*, and the completely phytochrome deficient *phyABCDE* mutant were insensitive to dose-dependent ABA application, implicating phy in ABA signalling (Yang et al. 2016). Interestingly, ABA application to WT effectively elicited rises in glucose, fructose, malate, fumarate and proline (Figure 4a, S8). This was also the case in *phyABD* and *phyABDE*, but here the ABA-induced changes are less pronounced. In untreated *phyABD* and *phyABDE*, proline levels exceeded those in WT plants treated with ABA, and did not alter with ABA application (Figure 4A). These data indicate that a common set of metabolites are regulated by phy and ABA, while the reduced ABA-sensitivity of *phyABD* and *phyABDE* suggests that phys partly operate through an ABA pathway (Wang et al. 2018).

**Figure 4:**
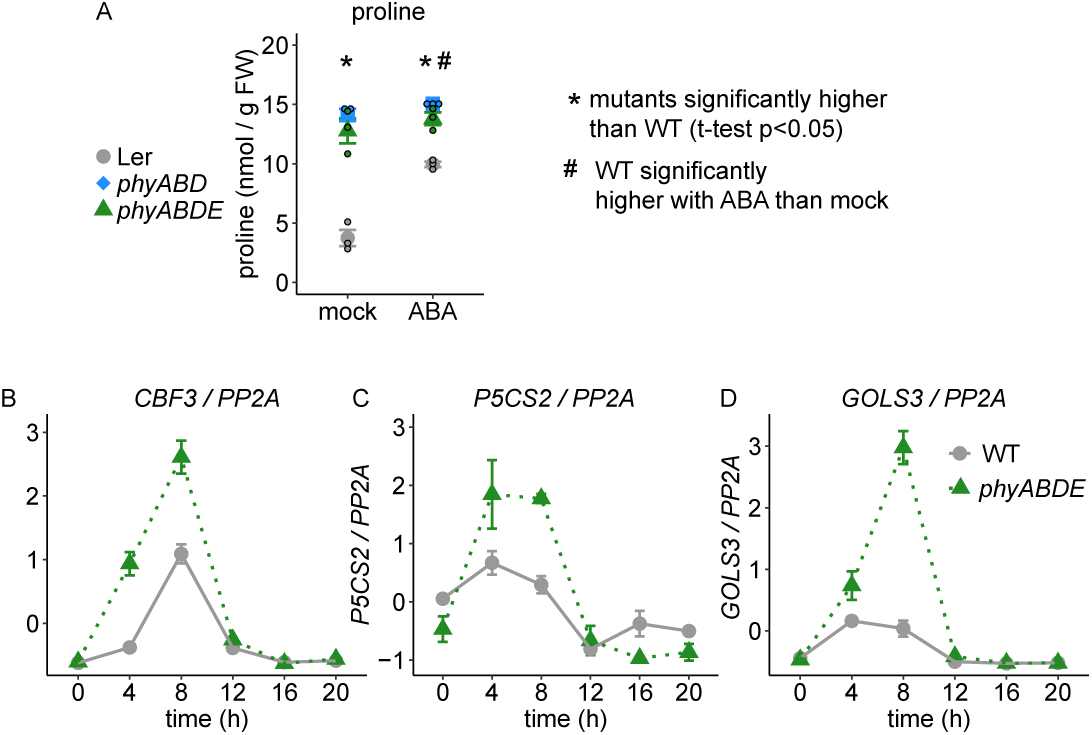
Testing involvement of abiotic stress signalling pathways. (A) Proline content in WT, *phyABD* and *phyABDE* at end of day after spraying twice a day for 2 days with either 100µM ABA, or a mock solution. (B-D) Transcript analysis of three cold response genes that have previously been implicated in metabolite over-accumulation in response to abiotic stresses: *CBF3* (B), *P5CS2* (C) and *GOLS3* (D). *PP2A* was used as a reference gene. Transcript data were scaled by subtracting the average of time courses of both genotypes and dividing by the standard deviation. Errorbars: SEM.

Phy and ABA signaling are known to converge through the interaction of PIF1 and ABI5 (Kim et al. 2016), and at *C-REPEAT BINDING FACTOR (*CBF)s (K. A. Franklin and Whitelam 2007) which operate in the cold acclimation pathway. In *phyABDE, w*e observe elevated daytime transcript levels of *CBF3*, which has previously been shown to have a prominent role in cold induced metabolite over-accumulation (Cook et al. 2004; S J Gilmour et al. 2000) (Fig. 4B). *phyABDE* mutants also exhibit elevated daytime expression of the CBF-regulated genes *DELTA1-PYRROLINE-5-CARBOXYLATE SYNTHASE* (*P5CS) 2* (Fig. 4C), which catalyzes a key step in proline biosynthesis, and *GALACTINOL SYNTHASE* (*GOLS) 3*, that catalyzes the first dedicated *de-novo* raffinose biosynthetic reaction (S J Gilmour et al. 2000; K. Maruyama et al. 2009; Taji et al. 2002). Collectively our data suggest that phy and ABA signalling may intersect in the regulation of metabolism, most notably proline synthesis.

### Phy-deficiency does not alter the rate of day-time protein and cell wall synthesis

As metabolite abundance reflects a balance between synthesis and utilization we wanted to establish whether usage of primary metabolites by major metabolite and energy consuming processes differed or was similar in phy-deficient mutants and WT. In growing cells, a large proportion of the free amino acid pool (Hildebrandt et al. 2015) and also indirectly the TCA cycle organic acid pool (Sweetlove et al. 2010) are used for protein synthesis. Thus, altered rates of protein synthesis could alter these metabolite pools. Here we used ^13^C incorporation methods to measure protein synthesis and, as a further indicator for C utilisation for growth, cell wall synthesis.

First we measured total protein content in plants of different ages and found that in older plants (30/35DAS) protein levels were similar in WT and *phyABD*. However, in younger plants (17-19DAS), protein levels were slightly lower at the start of the day (Fig 5A, fig S9A). Next, we measured protein synthesis in *phyABD* and *phyABDE* compared to WT by measuring the 13C incorporation into alanine in protein hydrolysate, correcting for incomplete labelling of the precursor by measuring enrichment in alanine in the free amino acid pool (Ishihara et al. 2015) (figure 5B, S9B). We found that rates of protein synthesis were very comparable between WT and mutant plants. Likewise, we found similar rates of ^13^C incorporation into cell wall cellulose in *phyABD, phyABDE* and WT (Figure 5C, S9C). Since water content is comparable in adult rosettes (Fig S1A), the amount of cell wall material is not expected to differ.

**Figure 5:**
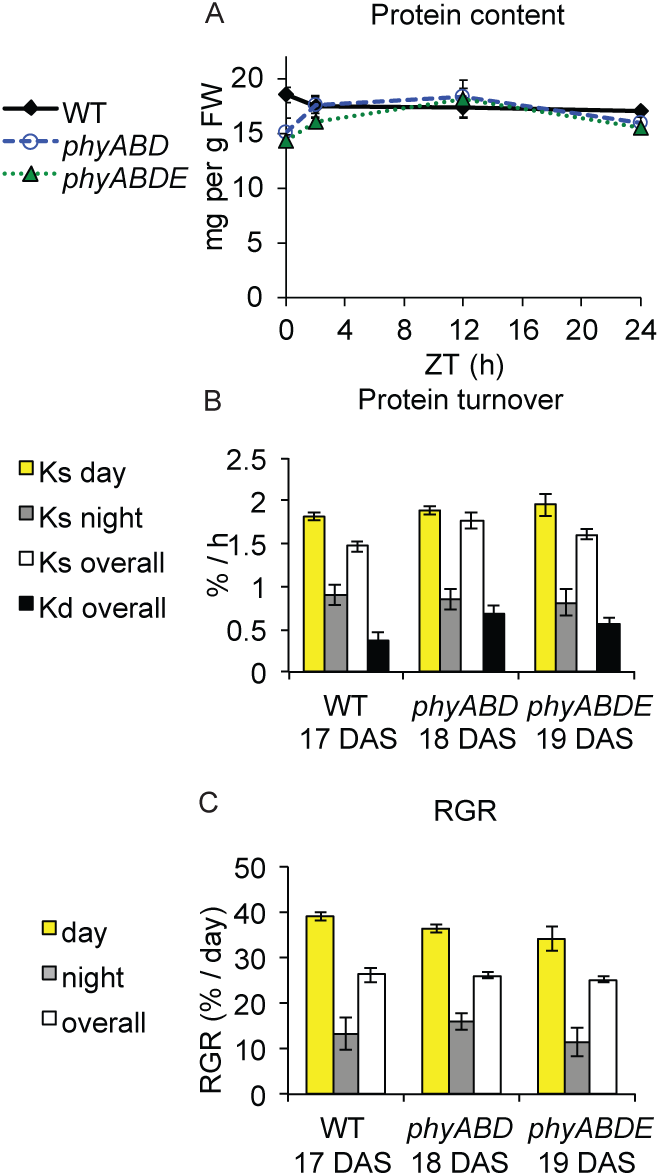
Protein turnover and relative growth rate (RGR) in ^13^C labelled samples in experiment 2. A) Protein content B) Protein synthetic rate (Ks) was calculated from incorporation of ^13^C into alanine in protein, adjusted by the ^13^C incorporation into free alanine. Degradation rates (Kd) were determined by subtracting RGR from Ks (see methods and Ishihara et al. 2015). C) RGR was calculated from ^13^C incorporation into cell wall cellulose. ^13^C incorporation at ZT12 was used for calculation of day time RGR, at ZT24 for overall RGR, and the difference between ZT12 and ZT24 for night time RGR. Error bars: (propagated) SEM. * p < 0.05.

The data therefore suggest that reduced consumption by protein and cellulose biosynthesis is unlikely to be the reason for over-accumulation of those metabolites with decreased synthetic rates. It should be noted that we cannot exclude a partial contribution of these biosynthetic processes as the total carbon content of the excess metabolites in *phy* mutants is small compared to the daily increase in dry biomass (Figure S10).

### The final biomass of *phy* mutants is determined at the seedling stage

Our published data show that one of the most striking features of severely phy-depleted plants is their reduced biomass, which in the case of *phyABDE* is only about 20% of WT plants (Yang et al. 2016). As we had not detected altered daytime ^13^C incorporation into protein and the cell wall in adult plants (Figure 5C, S9C) at this stage of the life history overall growth rate may be similar in *phy* mutants and WT. As already mentioned, phy-deficient seedlings had lower metabolites levels (Figure 1, Figure S1). We therefore reasoned that the final biomass phenotype could result from reduced growth during early development.

To test this hypothesis we first measured biomass and calculated the relative growth rate (RGR, = gain in biomass per day / pre-existing biomass) for different time intervals through development. Our data show that *phyABD* and *phyABDE* grow at a slower rate than WT until about 2 to 2.5 weeks (fig 6F). Thereafter, RGR is comparable to WT.

**Figure 6:**
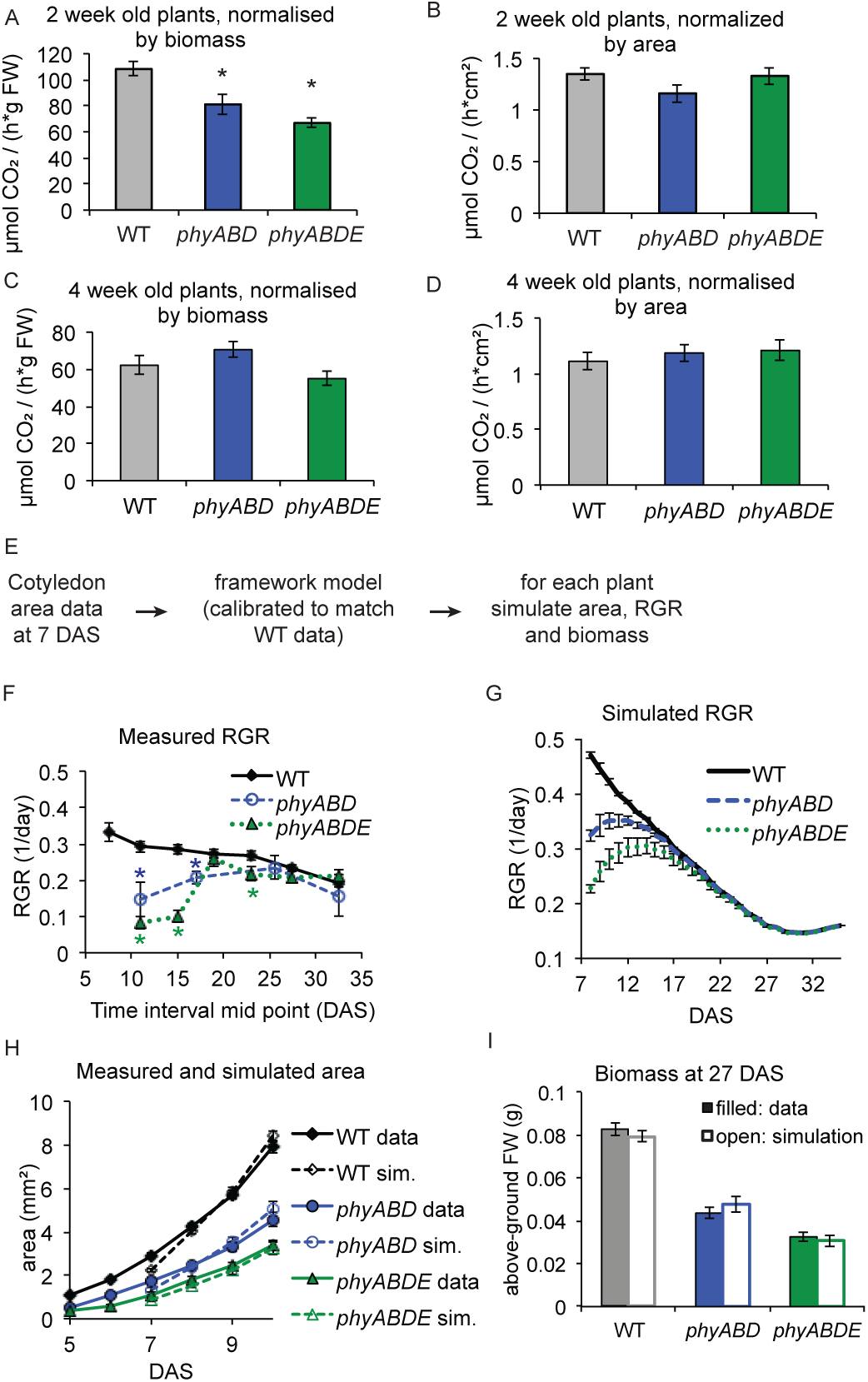
Final biomass deficit of *phy* kos is primarily due to reduction in RGR early in development, likely because of the reduced cotyledon size in *phy* knockouts. A-D) Net carbon uptake in 2 (A,B) or 4 (C,D) week old plants per unit biomass (A,C) or leaf area (B,D). E-I) Cotyledon data at emergence was used to predict RGR and final biomass using the *Arabidopsis* framework model. E) Simulations workflow F) destructive RGR time course measurement, G) simulated RGR timecourse of WT, *phyABD* and *phyABDE*. H) Measured cotyledon areas and simulated areas using 7 DAS cotyledon area data. I) measured and simulated biomass at 27 DAS. *p < 0.05 in t-test compared to WT

In agreement with our growth data, *phyABD* and *phyABDE* have a reduced rate of net carbon uptake per unit biomass at two but not at 4 weeks after sowing (figure 6A,C). Due to the decreased size of phy mutant cotyledons and first true leaves, when carbon uptake is normalized to cotyledon and leaf area instead of biomass, these differences disappeared (figure 6B,D). This suggested to us that at the fluence rates we had used, reduced leaf and cotyledon area available for photosynthesis in the *phy* mutants may be responsible for the reduced carbon uptake per unit biomass.

To test this hypothesis quantitatively, we used the *Arabidopsis* framework model. This model incorporates molecular mechanisms such as the circadian clock and photosynthesis but also carbon resources partitioning, organ formation and architecture, and can be used to simulate plant growth and development in different environmental conditions (Chew et al. 2014). In our experimental conditions, the model predicted that significant contribution of photosynthesis to growth started at 7 DAS. Therefore, we used individual measurements of cotyledon area of Ler, *phyABD* and *phyABDE* plants at EOD of day 7 as the starting cotyledon area in the model for each genotype. The model was calibrated for the WT to match 27 DAS biomass data by changing the input light intensity (supplementary methods). In good agreement with our RGR measurements the model predicts that *phy* mutants grow more slowly than the WT until about 2 to 2.5 weeks (figure 6F,G). The model simulation also correctly predicted phy mutant leaf area expansion rate during early development and final mature plant biomass. Leaf area and biomass measurements showed that model predictions were very accurate (figure 6H,I).

Taken together, our data show that reduced cotyledon area in phy-depleted seedlings restricts growth during early development, which severely compromises adult plant biomass.

### Accelerated seedling growth rescues the phyABD biomass defect

To definitively test whether delayed seedling growth can account for the severe reductions in *phyABD* adult plant biomass we identified a condition (long photoperiod, 18L:6D, 350µE light intensity) that accelerated early growth of *phyABD* to WT levels. WT and phyABD seedlings were grown in these condition until 10DAS, after which they were returned to our default 12L:12D, 115µE environment until sampling at 18DAS. Remarkably, this 10 day treatment was sufficient to completely rescue the *phyABD* adult biomass defect (figure S11A). This data therefore lends supports to the model prediction that slow growth early on can account for the adult biomass phenotype. Interestingly, even though biomass was restored to WT, *phyABD* still over-accumulated metabolites, suggesting that their accumulation occurs independently of any change in plant biomass-(figure S11B-D).

## Discussion

### *phy* mutants undergo a developmental switch in metabolism

It is well documented that phy-deficiency invokes a SAR, typified by leaf hyponasty, smaller leaf blades and longer petioles. We previously showed that these phenotypic changes are accompanied by an over-accumulation of sugars, amino acids and organic acids (Yang et al. 2016). Temporal tracking of malate, proline and starch revealed that these reference metabolites do not over-accumulate in *phyABD* and *phyABDE* at the seedling stage, rather, levels tend to be similar to WT or lower. Further, in *phyABD* and *phyABDE* seedlings, the balance between starch and glucose is shifted, with low starch and high glucose, when compared to WT. This suggests that carbon partitioning to starch vs sugars is altered in *phyABD* and *phyABDE* seedlings. A recent study analysing the starch biosynthesis mutants *pgm1, pgi1, agd2*, and the starch degradation mutant *sex1* showed that seedling hypocotyl elongation negatively correlates with starch levels (de Wit et al., 2018). Mutants with the longest hypocotyls had the lowest starch levels, suggesting that sugars that are not partitioned to starch are used to grow. The shift in the balance of starch and glucose may therefore contribute to the extreme elongated hypocotyl phenotype of *phyABD* and *phyABDE* seedlings.

We established that in adult phy-deficient plants, glucose, malate and proline over-accumulation is robustly observed over wide-ranging conditions, in petiole as well as blade tissue, and in the Col-0 and Ler accessions (Figure 1). In *phyABD* and *phyABDE*, metabolites begin to over-accumulate after 2 weeks, which coincides with the time mutant relative growth rate (RGR), which is initially slow, has adjusted to WT pace (Fig S1A, Fig 6).

### Phytochrome controls daytime levels of T6P

In agreement with our published work, LC-MS/MS and GC-MS data illustrate that phy depletion enhances TCA cycle intermediates, notably, fumarate, succinate, malate and isocitrate, and several amino acids of which proline and serine show the strongest changes (Yang et al. 2016).

LC-MS/MS enabled the detection of many phosphorylated and short-lived metabolites like pyruvate or glycerate (Arrivault et al. 2009; Szecowka et al. 2013). While many of these metabolites were not significantly altered, we established that daytime levels of T6P, Gal-1-P, Glc-1-P and Suc-6-P were elevated in *phyABD* (Fig 2, Fig S3). T6P typically parallels levels of sugars especially sucrose (John E Lunn et al. 2006; Yadav et al. 2014) and therefore its increased abundance in *phy* mutants may be a consequence of sugar over-accumulation.. It has been shown using inducible genetics that T6P stimulates flux to organic acids and amino acids, and that this involves post-translational activation of PEP carboxylase and nitrate reductase (Figueroa et al. 2016). Thus, the rise in T6P might explain part of the metabolic phenotype in the phy mutants, though not the large increases in proline and raffinose, which were not affected by inducible increases in T6P (Figueroa et al. 2016) and are more likely related to an abiotic stress response in phy mutants (see below). Interestingly, T6P has recently been shown to be an indirect modulator of PIF4 activity at higher temperatures (Hwang et al. 2019): KIN10, a catalytic subunit of the energy-sensing protein kinase SNF1-RELATED KINASE1 (SnRK1)complex was shown to phosphorylate and destabilize PIF4, preventing its accumulation at high temperatures. Genetic data suggest that T6P can indirectly promote PIF4 action by supressing the kinase activity of KIN10 (Hwang et al. 2019). As PIF4 levels and activity are likely to be elevated in phy-deficient plants (Lorrain et al. 2008; Johansson et al. 2014) high daytime levels of T6P may augment PIF4 action.

### Phytochrome suppresses stress metabolite synthesis in cooperation with ABA

We used a ^13^C labelling approach to determine whether *phy* mutants accumulate higher levels of sugars, amino acids and organic acids due to increased production or decreased downstream usage. In higher order *phy* mutants raffinose, myo-inositol, proline, serine, glutamine and pyroglutamic acid are synthesized at higher rates (figure 3C,D, figure S4 to 7). As myo-inositol is a precursor of raffinose, and the amino acids can all be synthesized from 2-oxoglutarate (Fig 3B) (Sengupta et al. 2015), this suggests that these two synthetic pathways are subject to phytochrome regulation. Interestingly, in contrast to most other TCA cycle intermediates, 2-oxoglutarate levels are not increased in *phy* mutants according to our LC-MS measurements, indicating that it may be increasingly used for the synthesis of these amino acids. The proline, serine and glutamine synthesis branch is connected to carbohydrate metabolism via short pathways and so their regulation is less energetically costly than other amino acids where synthesis and degradation requires multiple reaction steps (Hildebrandt 2018).

Proline and raffinose accumulation typically occurs in abiotic stress conditions, such as cold stress, high salinity and drought (Kaplan et al. 2004; Urano et al. 2009; Kempa et al. 2008; Pagter et al. 2017). Synthesis of glutamine from glutamate is key for nitrogen assimilation into organic molecules and high nitrogen assimilation rates are predicted to occur during cold stress (Hildebrandt 2018). Thus, phy depletion appears to elicit features of an abiotic stress response. Concurring with this notion, an earlier study showed that low R:FR treatment shade or *phy* mutations significantly improve freezing tolerance (K. A. Franklin and Whitelam 2007). Additionally, we provide evidence that the phy-dependent metabolic stress response involves ABA signalling, as the impact of ABA on metabolite levels is perturbed by phy-deficiency (Fig 4, S8). We also established that CBF3, a known ABA-regulated gene, has markedly elevated expression in *phyABDE* compared to WT (Fig 4). As *CBF3* over-expression induces metabolic changes with strong similarities to phy deficiency, in particular high proline accumulation, this suggests a potential molecular regulatory route for phy (S J Gilmour et al. 2000). We also found that *phyABDE* mutants exhibited elevated daytime expression of *P5CS2*and *GOLS3*, which are proline, and raffinose biosynthetic enzymes, respectively (Fig 4). Both enzymes are encoded by one or more other genes, but specifically these isoforms appear to typically get up-regulated by cold, more so than by drought, with *GOLS3* being regarded as a direct *CBF* target gene (Kyonoshin Maruyama et al. 2004; Taji et al. 2002; Fowler and Thomashow 2002; S J Gilmour et al. 2000; Sarah J Gilmour, Fowler, and Thomashow 2004).

Interestingly, increased ABA levels and high expression of ABA synthetic genes have been reported for the *prr975* mutant which also over-accumulates amino acids and organic acids (Fukushima et al. 2009). Since PIFs and PRRs mutually repress each other (Martìn et al. 2018), we speculate that ABA up-regulation is caused either by low PRR or high PIF levels.

In summary, our data is compatible with a model in which signals due to phy depletion converge with stress signaling, in particular ABA and CBFs, to control increased synthesis of typical stress metabolites, and potentially also other components.

### Phytochrome controls glucose and fructose synthesis and diurnal dynamics

A number of metabolites, including several amino acids, were mildly elevated by phy-deficiency, and for the majority we did not detect any sizeable changes in synthesis rates. In contrast, the sugars glucose and fructose accumulated to much higher levels in *phy* mutants in the later part of the light period. In the WT, synthesis of glucose and fructose from ZT0 is rapid, reaching steady-state levels by ZT2 and then declining through the night. At ZT0 *phy* mutants typically have slightly higher sugar levels than WT, initial label incorporation rates are slower, but glucose and fructose continue to rise, greatly exceeding WT levels during the later part of the day.

The results of our growth experiments and ^13^C incorporation into protein and cell wall did not give evidence for reduced usage of carbon by cell wall growth or protein biosynthesis. Therefore, the down-regulated processes responsible for the decreased usage have yet to be identified and, given the relatively small size of the sugar pools, may be rather minor Nevertheless, we have shown that phytochromes play a role in regulating the abundance of hexoses through the diurnal cycle. It is likely that the changes in sugar abundance affect sugar and energy signalling.

Phys may operate as part of a larger network to regulate sugar levels in leaves in different environmental conditions. Reducing sugar levels in the light period are higher in short than long photoperiod conditions (Mengin et al. 2017). This is at least partly due to a restriction of C utilization for growth in the light in short photoperiods (Mengin et al. 2017) and it has been proposed that this may be due to light signalling being more fully reversed at dawn after a long night than short night (Flis et al. 2016). As high reducing sugars were also found in the *elf3* mutant (Flis et al. 2019) (Flis et al., 2019), it is possible that Phy signalling interacts with outputs from the EC to control leaf sugar dynamics.

### Phy action at the seedling stage has consequences for adult plant biomass

Our measurements on ^13^C incorporation into protein and the cell wall surprisingly revealed that adult *phy* mutants do not have a reduced RGR, in spite of their strikingly different biomass and leaf architecture. As seedlings severe *phy* mutants have low levels of starch and other metabolites (Fig 1C), we therefore reasoned that the dramatic biomass reductions observed in higher order *phy* mutants at the adult stage may be a consequence of delayed early development. We established, by quantifying biomass, that RGR was indeed slower in *phyABD* and *phyABDE* compared to WT during early development, but not in adult plants (Fig 6). Previous studies showed that higher order *phy* mutant seedlings have lower chlorophyll levels (Strasser et al. 2010; Hu et al. 2013). In agreement with these studies, and correlating with our growth data, we demonstrated that *phyABD* and *phyABDE* have a reduced rate of net CO_2_ uptake per unit biomass in seedlings, but not later in development (figure 6A,C). We also established that if carbon uptake is normalized to leaf area instead of biomass, the differences between WT and *phy* mutants diminish (figure 6B,D). As *phyABD* and *phyABDE* have small cotyledons, it is therefore possible that reduced area available for photosynthesis may be an important factor in limiting carbon uptake.

To formally test this idea we used the modular framework model, which provides a means to test the impact of impaired cotyledon expansion on photosynthesis, and plant biomass accumulation (Chew et al. 2014). Remarkably, by altering starting cotyledon parameters alone, the model could simulate cotyledon area expansion, relative growth rate through development and final plant biomass of WT, *phyABD* and *phyABDE* to a high degree of accuracy (Fig. 6). Thus, Framework model predictions strongly support our premise that final biomass reduction of *phy* mutants can be explained exclusively by the smaller cotyledon area at seedling emergence. Further corroboration comes from our data showing that a growth accelerating treatment applied at the seedling stage fully rescued the *phyABD* biomass defect. Our study establishes that the important role that phytochromes play in seedling establishment is critical for setting the pace of growth and ultimately adult plant biomass.

## Methods

### Plant material and growth conditions

The *phy* mutants (*phyBD* (Devlin et al. 1999), *phyABD* (Devlin et al. 1999), *phyABDE* (K. a Franklin et al. 2003) used in this study were all in the Ler background unless otherwise indicated. All mutants used have been previously described (*phyABD* mutant in Col0 (Sánchez-Lamas, Lorenzo, and Cerdán 2016), 35S:PIF4-HA and 35S:PIF5-HA (Lorrain et al. 2008), *hy5-215* (Oyama, Shimura, and Okada 1997).

Seeds were surface sterilised with 30% thin bleach and 0.01% TritonX-100, washed 5 times with sterile water and stratified at 4°C for 4 to 5 days in ddH2O. For gas exchange measurements and the 5 week growth curve analysis, where plants were grown entirely on soil, surface sterilisation was skipped, but 200µM GA4+7 (Duchefa) was added to the water during stratification and then washed off before placing on soil to improve germination of the higher order *phy* mutants (Sánchez-Lamas, Lorenzo, and Cerdán 2016).

With the exception of gas exchange experiments and the growth curves, stratified seeds were placed on plates (1/2 MS pH5.8, 1.2% agar), and grown at 18°C and 115µE either in LD8:16 photoperiods for experiments with 5 week old plants, or in LD12:12 photoperiods for experiments with younger plants. Exceptions to these conditions were for the experiments testing the conditionality of metabolite over-accumulation (Fig S2, supplementary methods), and the growth curve experiment was carried out in LD12:12 entirely. The same-biomass WT control in labelling experiment 1 and the LC-MS experiment was also grown at LD8:16 for 2 weeks and then at LD12:12 until harvesting at 30DAS.

Seedlings were transferred to soil at 14 (for 5 week old plant experiments) or 10 DAS (for younger plant experiments), 7DAS (labelling experiment 2 and 3) or 5 DAS (metabolite measurements in 10 and 14 day old seedlings). Growth conditions were kept the same after transfer to soil except for experiments on 5 week old plants where the light regime was switched to LD12:12.

Plants were sampled at the indicated developmental times and times of the day. In labelling experiments 2 and 3, plating and transfer of WT, *phyABD* and *phyABDE* had to be staggered by 1 day such that they were sampled at 17 DAS or 18 DAS or 19 DAS since transfer took too long to complete all three genotypes in one day, but still to allow labelling at the same day. These two were identical except that different seed batches were used.

When plants were sampled at the end of the day (EOD), this was done within the last 30min of the light period. Sampling was done in three replicates with at least 5 plants per replicate. Tissue was quickly cut and flash-frozen in eppendorf tubes in liquid nitrogen. In the case of plants for LC-MS analysis of phosphorylated and short half life metabolites (Fig 2c), rosettes were flash frozen in a mortar inside the incubator without changing the light exposure of the cut before freezing (Arrivault et al. 2009; Szecowka et al. 2013).

### ^13^C labelling

^13^C labelling with air containing only ^13^CO_2_ instead of ^12^CO_2_ was carried out as described by (Ishihara et al. 2015). ^13^C labelling was started at light onset and carried out for 24h. Samples were taken in triplicates, with 5 plants per replicate for experiment 1, 25 to 30 plants for experiments 2 and 3, at ZT0 (unlabelled), ZT2, ZT12 and ZT24.

### ABA treatment

ABA stock solutions of ABA was prepared in 100% DMSO at 50mM (ABA)The final sprayed solution contained 0.01% Silwet and 1.2% DMSO. 3 week old plants were sprayed at the beginning and the end of the day for 2 days, at the beginning and middle of the day on the third day and sampled at the end of the third day of spraying.

### Enzymatic metabolite measurements

Without allowing tissue to thaw, samples was crushed with a metal ball in eppendorf tubes using a Qiagen TissueLyzer and aliquots of about 20mg were weighed out and the aliquot weight was recorded for normalisation of metabolite abundance. All soluble metabolites (glucose, fructose, malate, fumarate, proline) were measured from ethanolic extracts as described by (Mengin et al. 2017). Glucose units from starch was measured after hydrolysis of the ethanol insoluble pellet after ethanolic extraction as described as in (Mengin et al. 2017). Protein content for Fig 4a-b was carried out by a standard Bradford assay.

### Gene expression analysis

For qRT-PCR analysis, three weeks old plants were used. Total RNA was isolated from approx. 70mg of finely ground tissue using Qiagen’s RNeasy Plant Mini Kit with on-column RNAase-free DNase digestion. cDNA synthesis was performed using the qScript cDNA Supermix (Quanta Biosciences) as described by the manufacturer. The qRT-PCR was set up as a 10μL reaction using SYBR Green (Roche) in a 384-well plate, performed with a Lightcycler-480 system (Roche). Results were analyzed using the Light Cycler-480 software. Expression values of target genes were normalized to the values of PP2A reference gene. The primers used in this study are given in Supplementary Table 1.

### Sample preparation for LC-MS/MS analysis

For LC-MS analysis of metabolite abundance at ZT6 and ZT24, as well as for analysing label incorporation into S6P, metabolites were extracted from ∼15mg tissue aliquots. Measurements and data analysis were carried out as described by (Arrivault et al. 2009).

### Sample preparation for GC-MS analysis

Soluble metabolites were extracted from ∼30mg aliquots with methanol, followed by a phase separation with chloroform and water as described previously (Arrivault et al. 2009; Ishihara et al. 2015).

Protein was extracted from the methanol-insoluble pellet. 50µg protein was precipitated with TCA, washed with acetone and subsequently hydrolysed and neutralised as described by (Ishihara et al. 2015).

After removal of protein from the sample pellet, starch was degraded, cell wall material was hydrolysed and neutralised, as in (Ishihara et al. 2015).

### Derivatisation and GC-MS analysis

After drying soluble metabolites or neutralised hydrolysates, samples were derivatised and prepared for GC-MS and mass spectrometrically analysed as described by (Lisec et al. 2006).

### Analysis of GC-MS data

The XCalibur™ software (Thermo Fisher Scientific Inc., version 2.2 SP1.48, 2011) was used for identification and quantification of metabolite peaks. cdf files were converted to .raw files and imported into the Processing Setup tool. Peaks were manually assigned by using standards run just prior to the samples, retention index and fragmentation spectra from the metabolite library (‘110524_modified_TFLIB (Version_20070220_05)_AFE_FEMS-Martin_C13_Sorbitol.xls’ from the Golm metabolite database). The XCalibur™ Sequence batch processing tool was used to identify the metabolites of interest in all samples. Peak selection was manually verified and adjusted were needed, using the XCalibur™ QuanBrowser tool. The metabolite analyte IDs and masses used are included in supplementary dataset 1

The resulting peak abundance data of each dataset was analysed, resulting in metabolite abundance as well as 13C enrichment for each metabolites. Results were normalised by the ribitol abundance as well as aliquot FW. The enrichment data was adjusted for natural occurrence of different carbon isotopes using the corrector software version 10 (Huege et al. 2014). The latest time point before saturation of label incorporation was determined for each metabolite, and the corresponding % label incorporation was multiplied by the abundance at the same time point to obtain a measure for the relative incorporation of ^13^C label carbon in this time period. Relative abundance and relative incorporation of new carbon were reported in Figures S4-7 and ratios of phy mutant / WT were computed for heatmaps. T-tests were used to determine significant differences between WT and phy mutants.

### RGR and protein turnover calculations

RGR and protein turnover were calculated as described by (Ishihara et al. 2015). For daytime overall RGR, 13C incorporation at ZT12 and ZT24 was used, respectively. For night-time RGR, incorporation at ZT12 was subtracted from incorporation at ZT24, with the SEM of each time point used for error propagation. For Ks values, incorporation of 13C into alanine in protein hydrolysate was used for the same time points as for RGR. To adjust for differences in free alanine labelling, 13C of free alanine at ZT12 was used for day-time Ks, at ZT24 for overall Ks, and the average of ZT12 and ZT24 for night-time Ks.

### Photosynthetic gas exchange measurements

For measuring net carbon uptake in 2-week old plants, grown in L12:12, 18°C and 115µE, a multi-chamber system as described by (Kölling et al. 2015) together with a LiCOR-7000 CO2/H2O was used. Each replicate consisted of a pot with at least 20 plants on soil, and 4 replicates were measured for each genotype. Gas exchange was measured after stabilisation in the chambers for 30min at 115µE. Plants were photographed just before measurement for determination of leaf area, and immediately cut off after the measurement to determine total above-ground biomass. Pots with soil were then placed back into the photosynthesis chamber to determine background gas exchange by the soil, which was subtracted from the measurement with plants.

For measurement of 4 week old plants, grown in the same conditions as the 2 week old plants, the setup described by (Mengin et al. 2017) was used (LI-6400XT Portable Photosynthesis System with a whole-plant Arabidopsis chamber and a 6400-18 RGB light source), with a light intensity of 115µE and 60% humidity and 2 or 3 plants per pot.

### Simulation of RGR, biomass and rosette area using the *Arabidopsis* framework model

Cotyledon area was monitored at EOD from 5 DAS to 10 DAS in WT, *phyABD* and *phyABDE* seedlings growing on soil, 110µE, 18°C and LD 12:12. Cotyledon area was analysed using Adobe Photoshop. Plants were transplanted to individual pots after the imaging and were grown in the same conditions until 27 DAS when their above-ground biomass was measured.

The framework model was calibrated by changing the light intensity input to correctly predict the WT 27 DAS biomass from the WT cotyledon area at 7 DAS, which is the day the model predicted emergence under the conditions used. Subsequently, the calibrated model was used to simulate cotyledon area, RGR and biomass for all three genotypes, setting the parameter em to the measured cotyledon sizes.

## Supplementary tables

**Supplementary table 1:**
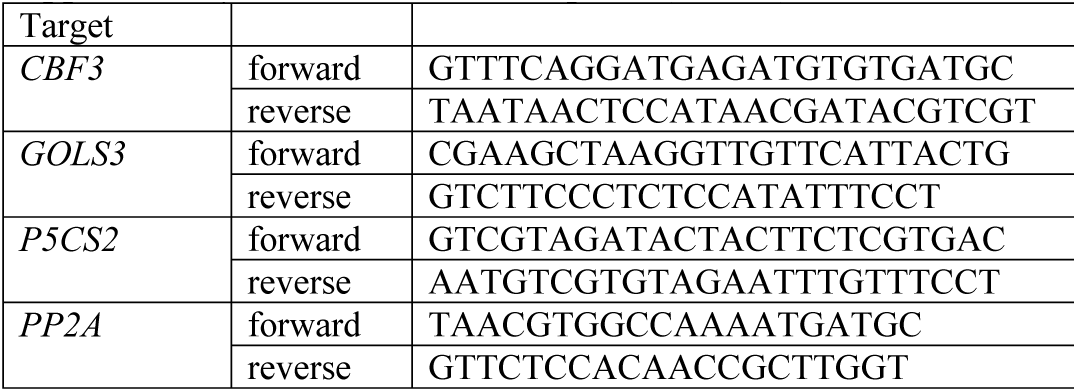
Primers for qPCR (5’ to 3’)

## Supplementary data

Data S1: All GC-MS and LCMS data.

## Supplementary methods

### Growth conditions for testing conditionality of metabolite over-accumulation

Our reference conditions for these experiments were as described for 5 week old plants in the methods section in the main test.

For measurement in different photoperiods (Fig S2A-C), plants were either kept at LD8:16 after the initial 2 weeks in short days, or moved to either LD12:12 or LD16:8 for the remaining 3 weeks. To test the effect of light intensity (Fig S2D-F), plants were grown in our reference conditions but were moved to different light intensities for the last 3 days before sampling at the end of the day. For temperature experiments (Fig S2G-I), plants were moved to either 16°C or 22°C 4 days prior to sampling.

### Water content measurement

For water content measurements (Fig S1A), fresh weight (FW) of groups of seedlings was determined which were then dried at 80°C for 3 days and weighed again for dry weight (DW) and water content was calculated ((FW – DW) / FW * 100%).

---

**Supplementary figure S1:**
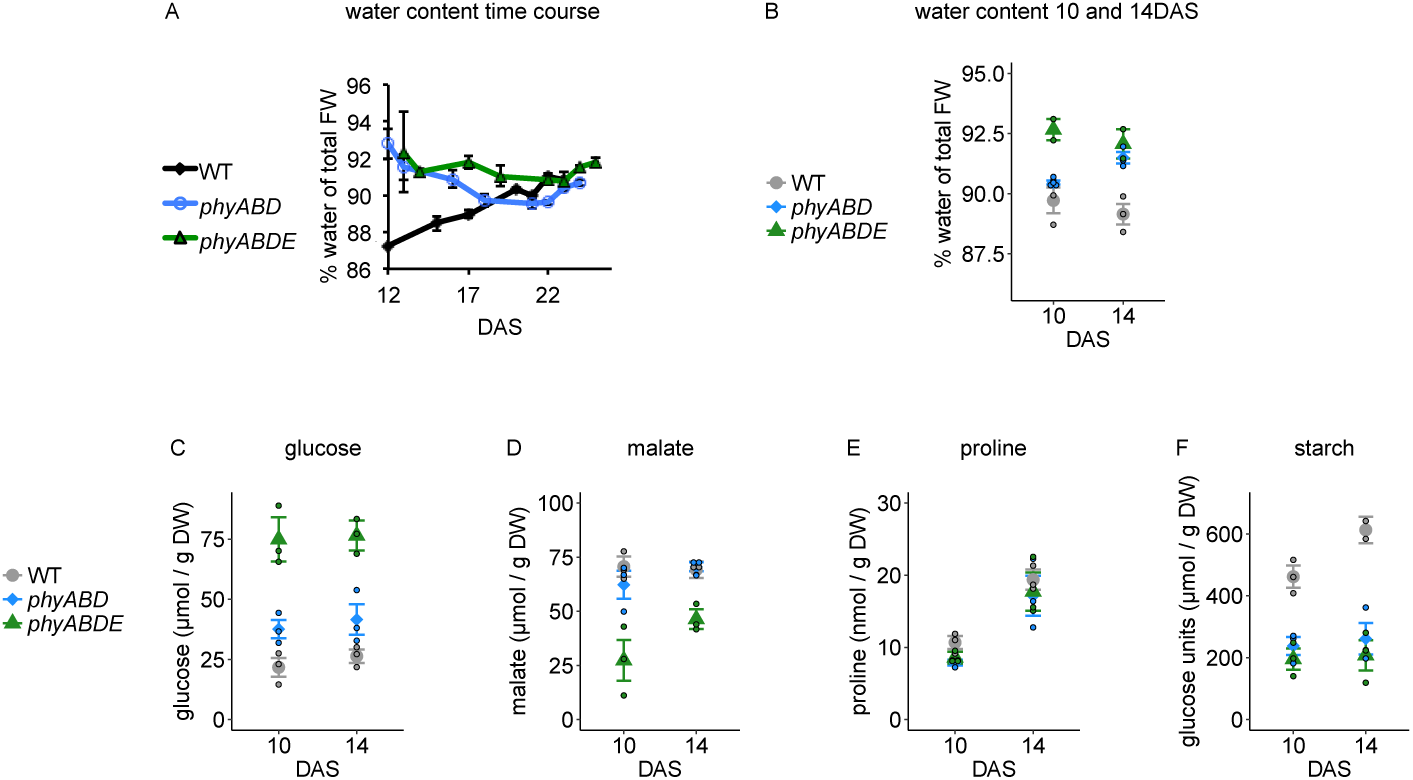
Water content in WT and *phy* mutants and metabolites in seedlings normalised by DW. A) Water content over a developmental time course was measured in WT, *phyABD* and *phyABDE*. B) Water content at 10 and 14 DAS of seedlings grow in parallel to those in figure 1. C-F) Metabolite content of experiment shown in figure 1 but normalised by DW using data in (B). C) glucose, D) malate, E) proline, F) starch. Legend symbols indicated mean, errorbars SEM, coloured circles represent the individual measurements of replicates.

**Supplementary figure S2:**
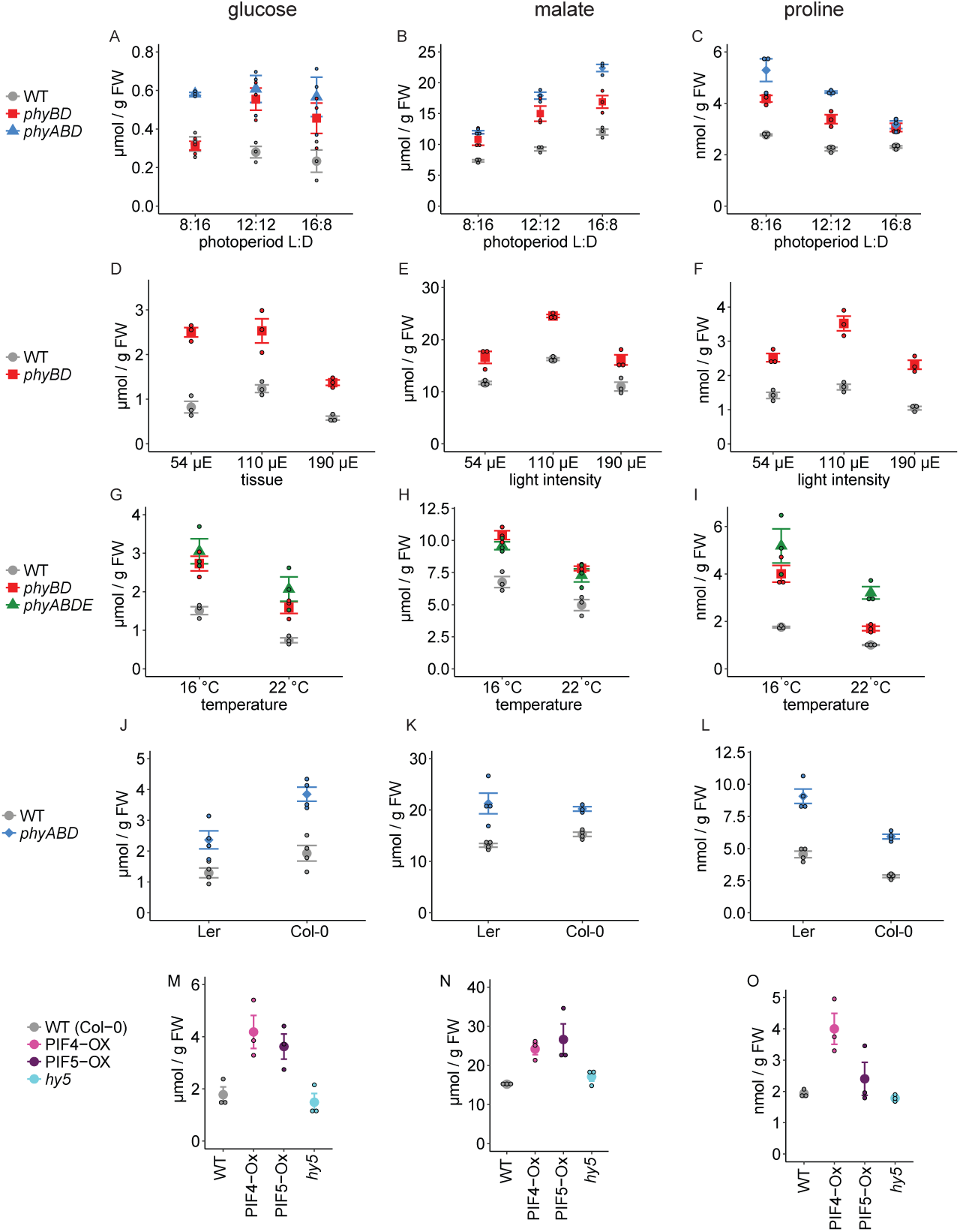
Metabolite content in *phy* mutants and WT in different conditions and genetic backgrounds. Glucose (A,D,G,J,M), malate (B,E,H,K,N) and proline (C,F,I,L,O) were measured in 5 week old WT and mutants. Unless otherwise indicated, conditions were 115 µE, 18°C, LD 8:16 for 2 weeks, followed by LD 12:12 for 3 weeks. (A-C) Plants were shifted to different light intensities (54 µE, 115 µE or 190 µE) 3 days before sampling. (D-F) Plants were grown in different photoperiods (LD 8:16, LD 12:12 or LD 18:8) after the initial 2 weeks in short days. (G-I) Plants were shifted to different temperatures (16 °C or 22 °C) 4 days before sampling. (J-L) Metabolites in the *phyABD* mutant in the Ler and Col-0 backgrounds with the corresponding WT controls. (M-O) Metabolites in WT, 35S:PIF4-HA, 35S:PIF5-HA and *hy5*.

**Supplementary figure S3:**
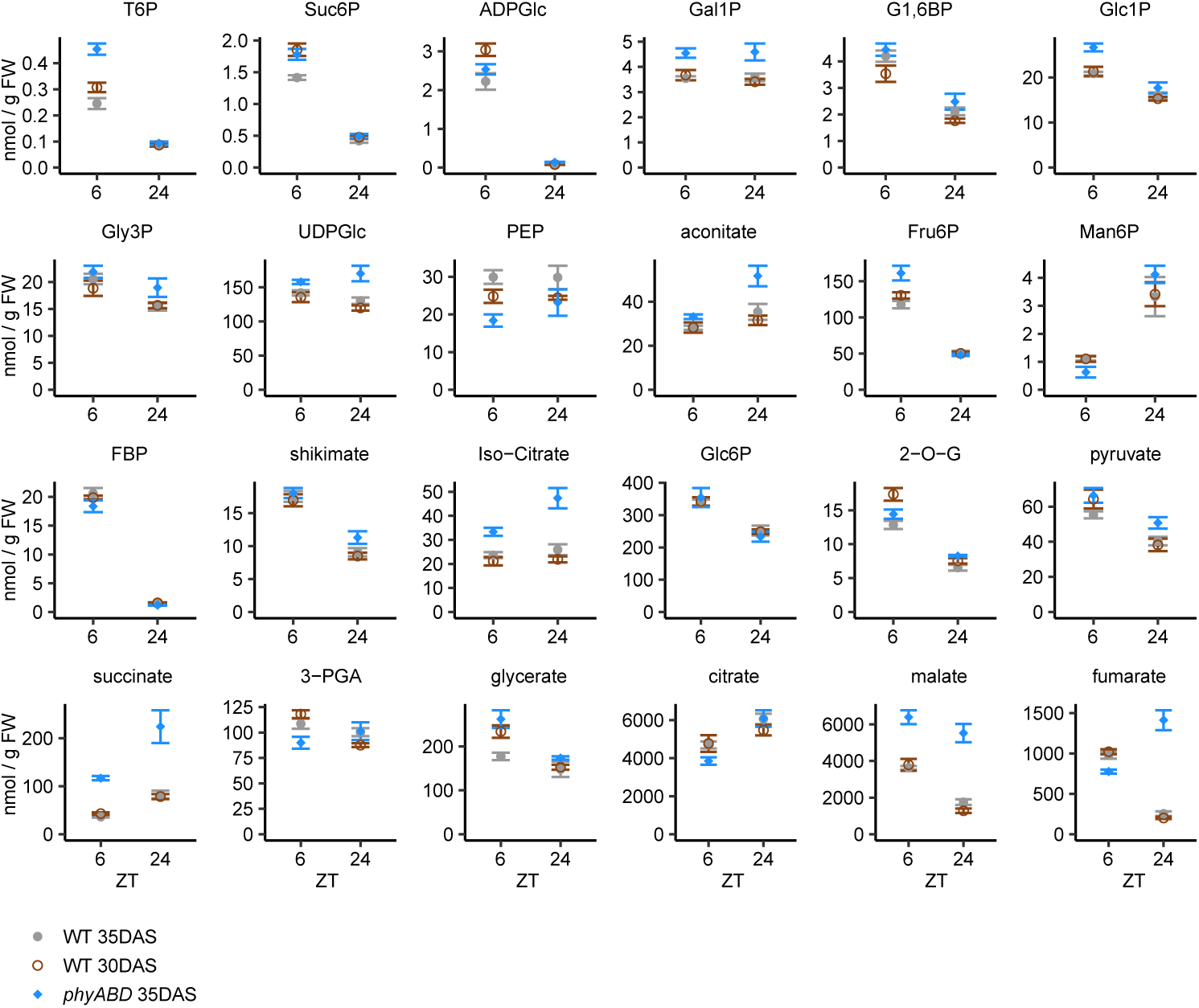
Metabolite abundance in WT and *phy* mutants measured by LC-MS/MS. A selection of metabolites was measured by LC-MS as it is better suited for analysis of phosphorylated and instable metabolites. A typical day time points (6h) and an end of night time point (24h) were selected for analysis of metabolites in 5 week old WT and *phyABD* plants as well as a same biomass WT control (30 DAS).

**Supplementary Figure S4:**
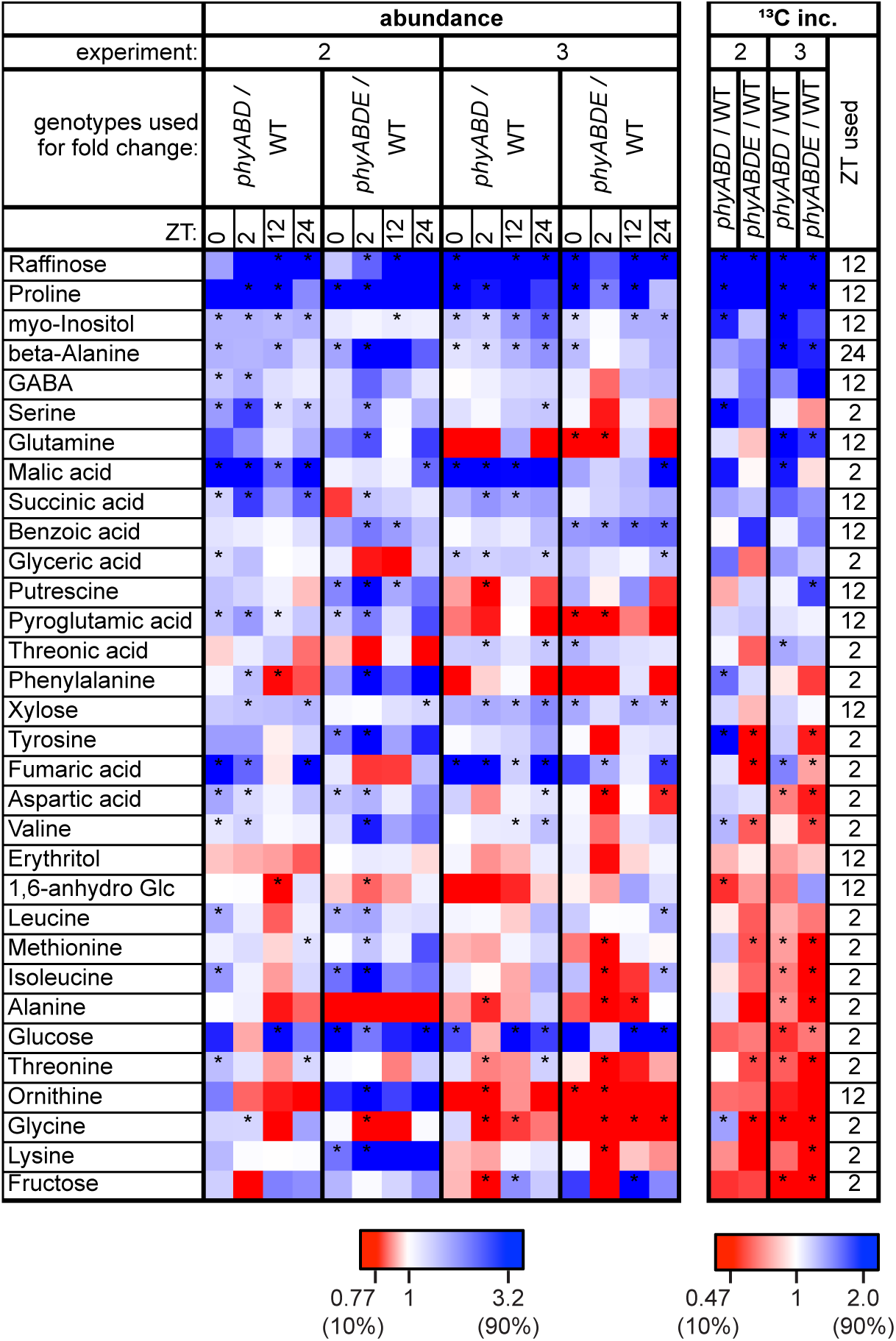
Metabolite abundance and label incorporation fold changes in *phy* mutants / WT in labelling experiments 2 and 3 (17 to 19 DAS). Heatmap displays fold change in abundance (left) and ^13^C label incorporation rate (right, ‘^13^C inc.’) of *phy* mutant value / respective WT control. All abundance time points are shown, but only most suitable time point for the incorporation rates (time interval indicated in the rightmost column). Metabolites are sorted by decreasing average fold change of incorporation rates.

**Supplementary figure S5:**
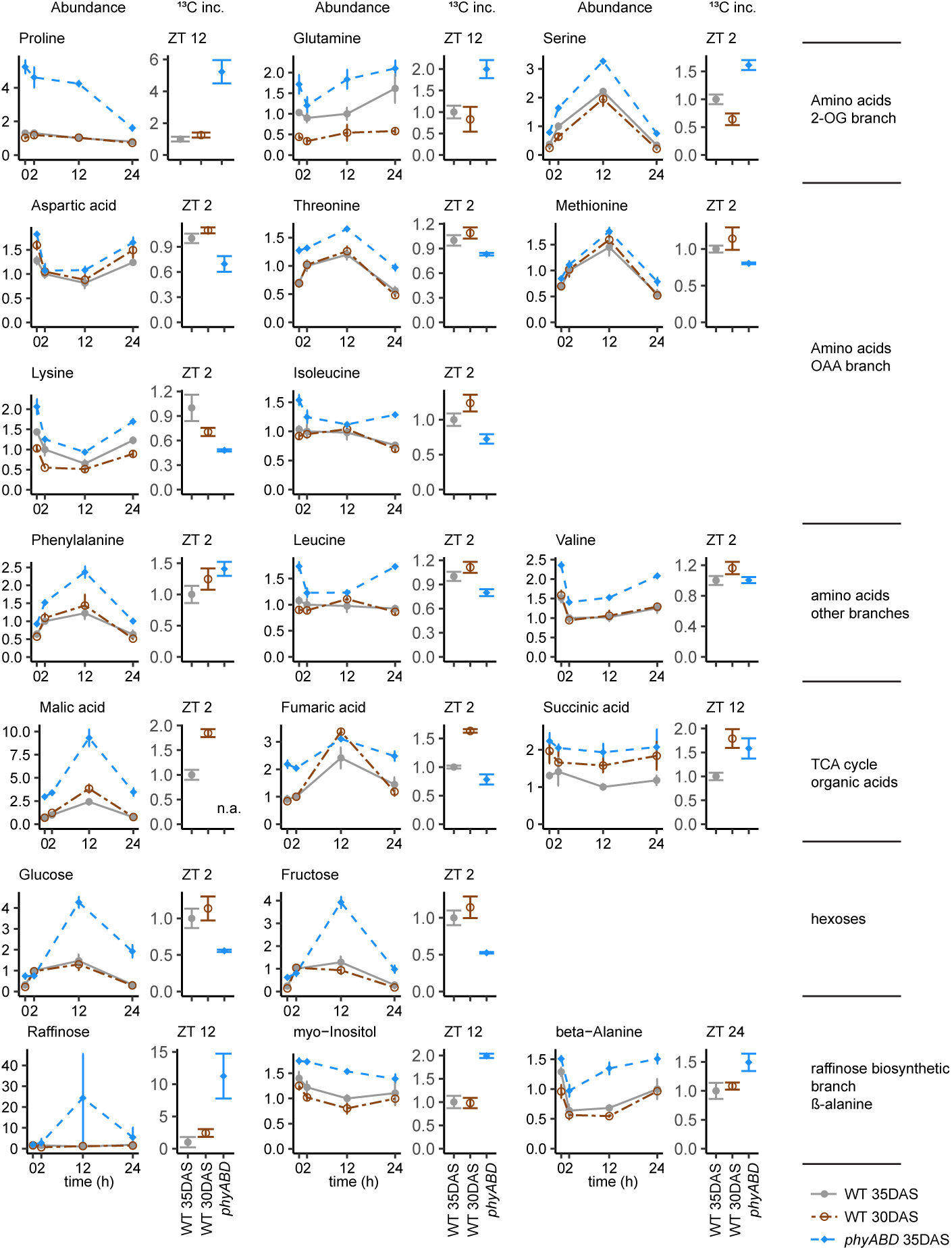
Plots showing GC-MS abundance and 13C label incorporation rate (‘^13^C inc.’) data of selected metabolites, labelling experiment 1. For each metabolite, abundance data is shown as a time course, the label incorporation rate, calculated by multiplying abundance and % ^13^C enrichment for the same time point, is shown for the most suitable time point, which is indicated above the label incorporation rate plots. Abundance and label incorporation rates were normalised by the 35DAS WT mean of the time point used for the label incorporation rate. Errorbars: SEM (abundance) or propagated SEM (label incorporation). n.a.: enrichment data for malate could not be determined.

**Supplementary figure S6:**
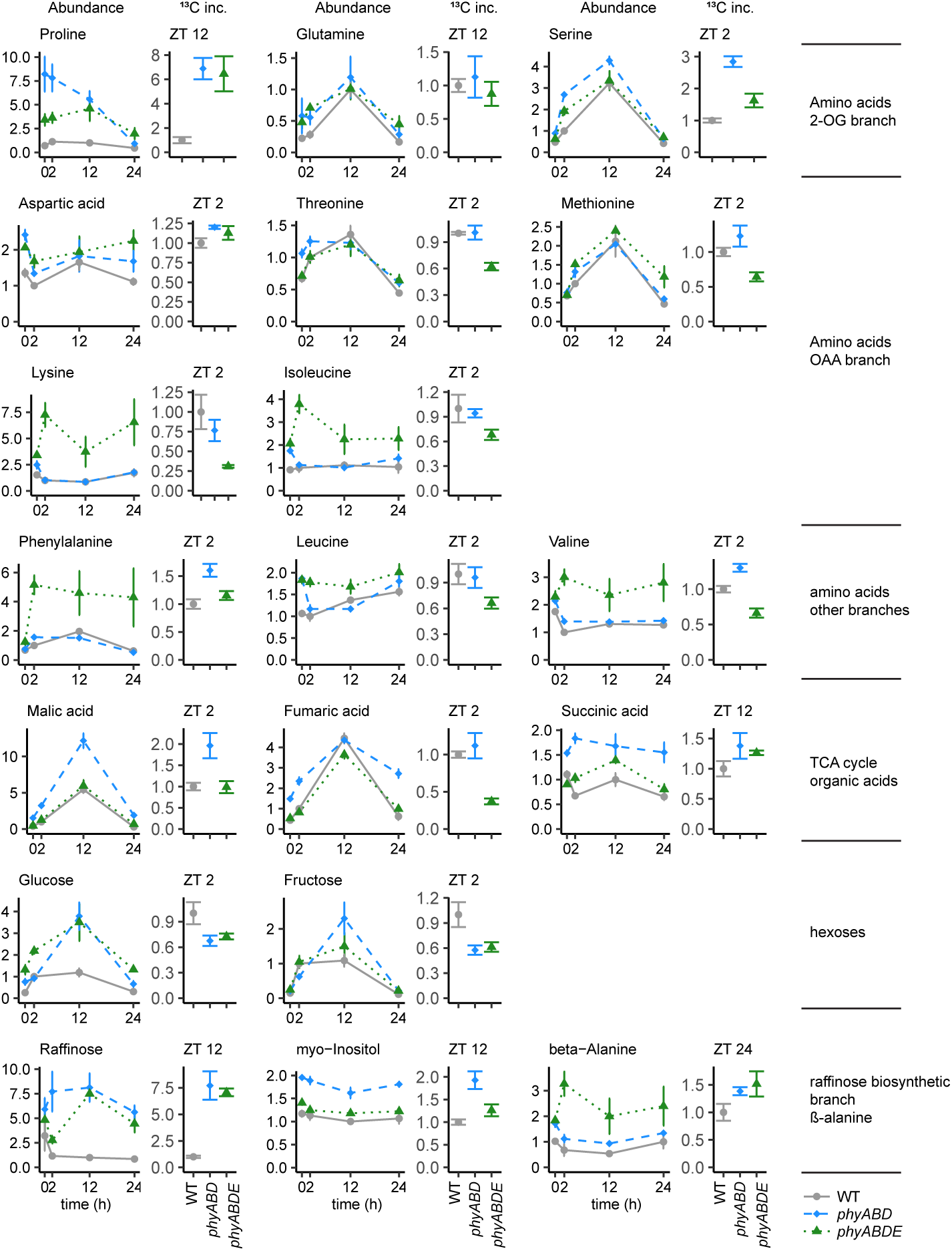
Plots showing GC-MS abundance and 13C label incorporation rate (‘^13^C inc.’) data of selected metabolites, labelling experiment 2. For each metabolite, abundance data is shown as a time course, the label incorporation rate, calculated by multiplying abundance and % ^13^C enrichment for the same time point, is shown for the most suitable time point, which is indicated above the label incorporation rate plots. Abundance and label incorporation rates were normalised by the 35DAS WT mean of the time point used for the label incorporation rate. Errorbars: SEM (abundance) or propagated SEM (label incorporation).

**Supplementary figure S7:**
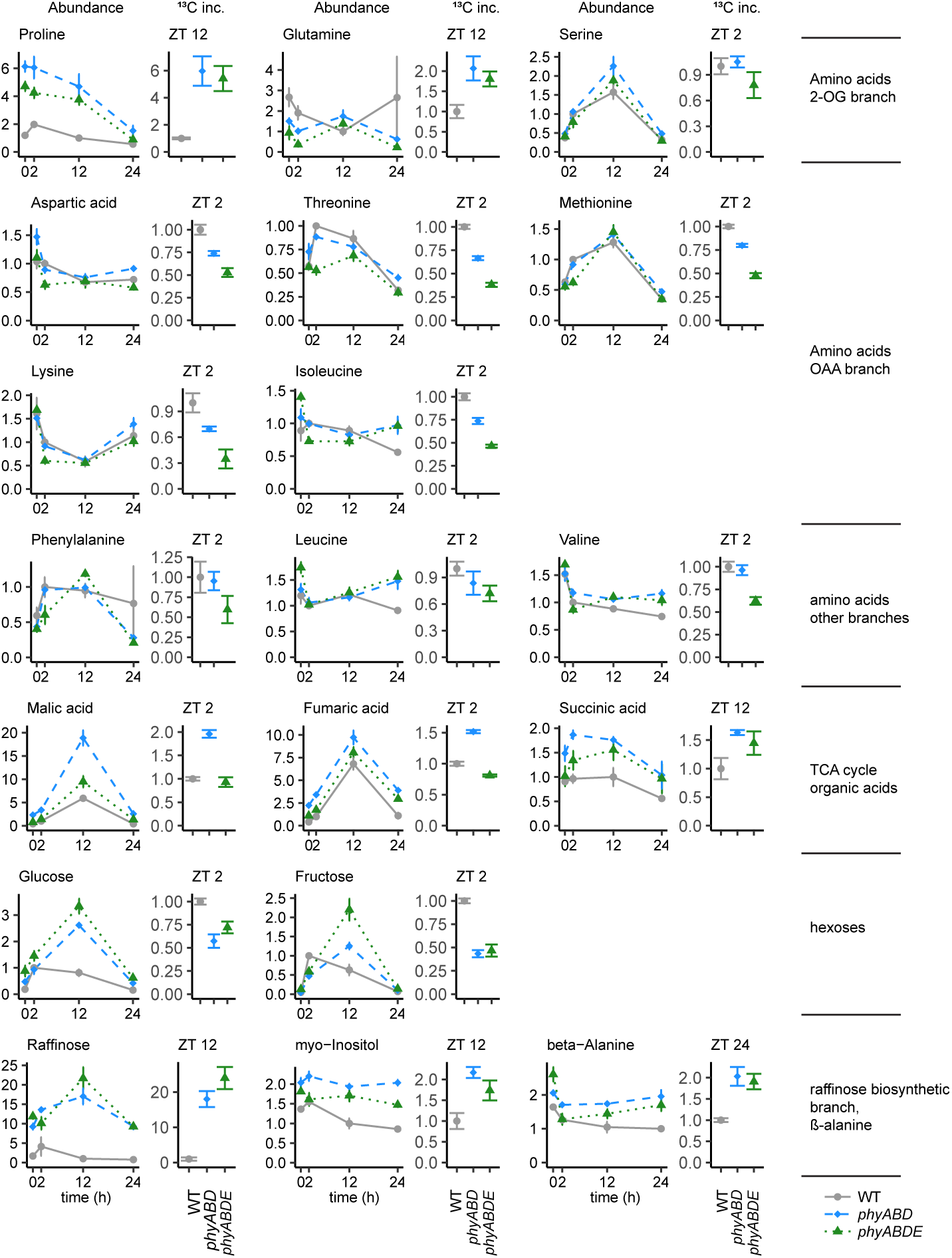
Plots showing GC-MS abundance and 13C label incorporation rate (‘^13^C inc.’) data of selected metabolites, labelling experiment 3. For each metabolite, abundance data is shown as a time course, the label incorporation rate, calculated by multiplying abundance and % ^13^C enrichment for the same time point, is shown for the most suitable time point, which is indicated above the incorporation rate plots. Abundance and label incorporation rates were normalised by the 35DAS WT mean of the time point used for the label incorporation rate. Errorbars: SEM (abundance) or propagated SEM (label incorporation).

**Supplementary figure 8:**
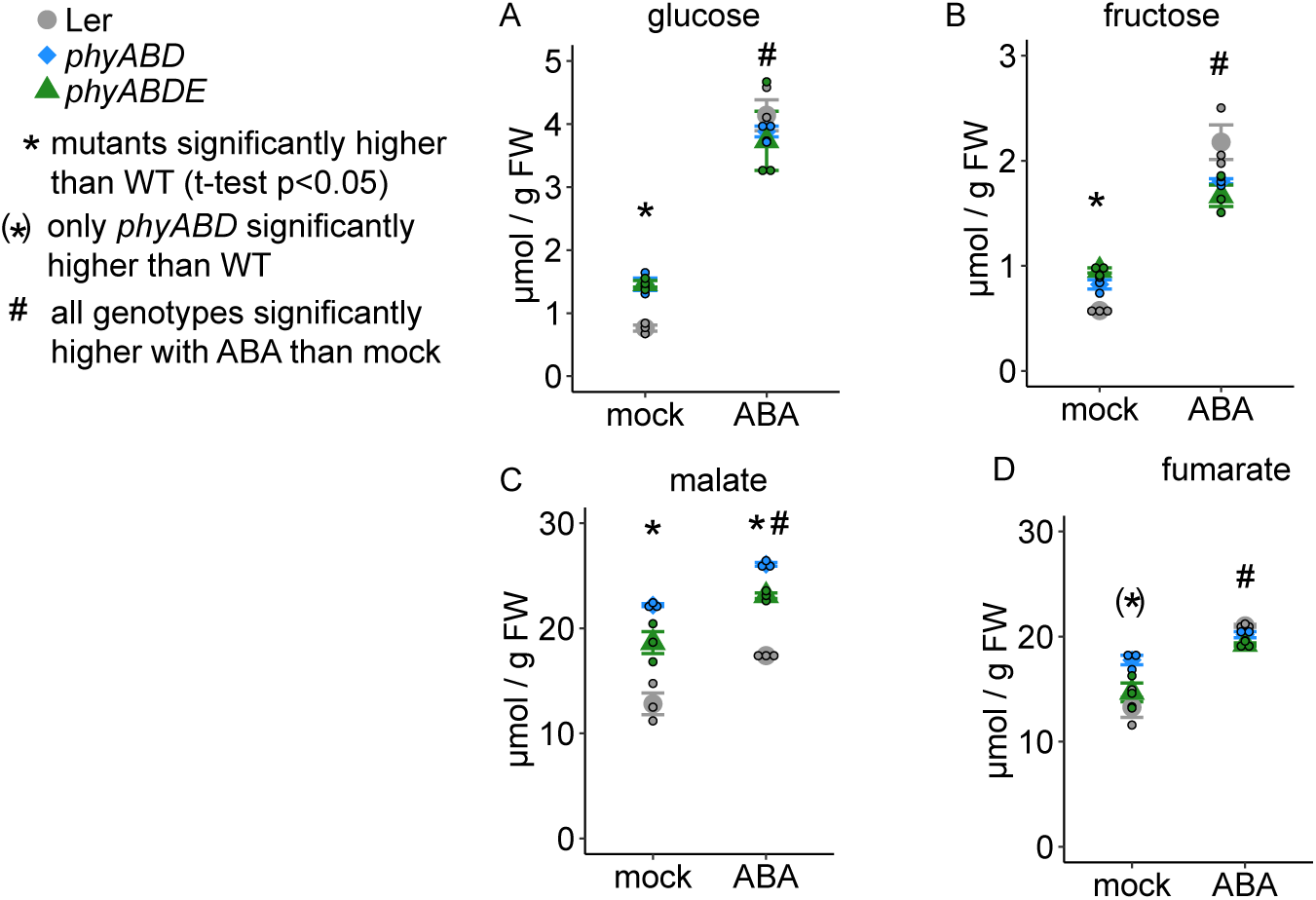
Effect of ABA treatment on glucose, fructose, malate and fumarate in WT and *phy* mutants. Metabolite content in WT, *phyABD* and *phyABDE* at end of day after spraying twice a day for 2 days with either 100µM ABA, or a mock solution. (A) glucose, (B) fructose, (C) malate, (D) fumarate. Errorbars: SEM. Little circles indicate individual replicate measurements.

**Supplementary figure 9:**
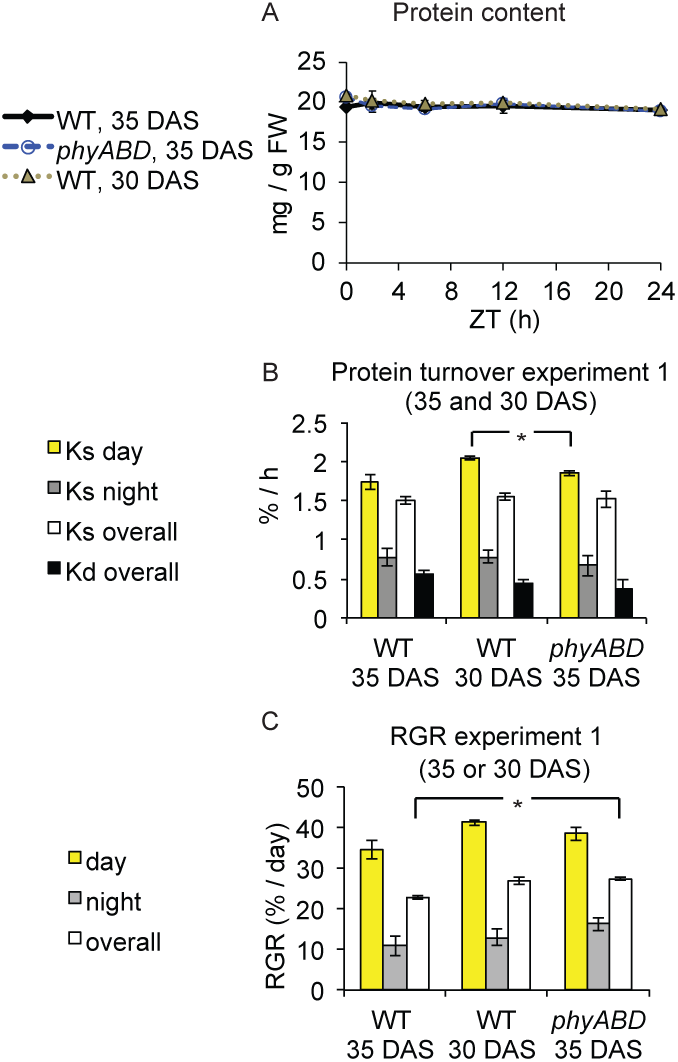
Protein turnover and relative growth rate (RGR) in ^13^C labelled samples in experiment 1. A) Protein content B) protein synthetic rate (Ks) was calculated from incorporation of ^13^C into alanine in protein, adjusted by the ^13^C incorporation into free alanine. Degradation rates (Kd) were determined by subtracting RGR from Ks (see methods and Ishihara et al. 2015). C) RGR was calculated from ^13^C incorporation into cell wall cellulose. ^13^C incorporation at ZT12 was used for calculation of day time RGR, at ZT24 for overall RGR, and the difference between ZT12 and ZT24 for night time RGR. Error bars: (propagated) SEM. * p < 0.05.

**Supplementary figure 10:**
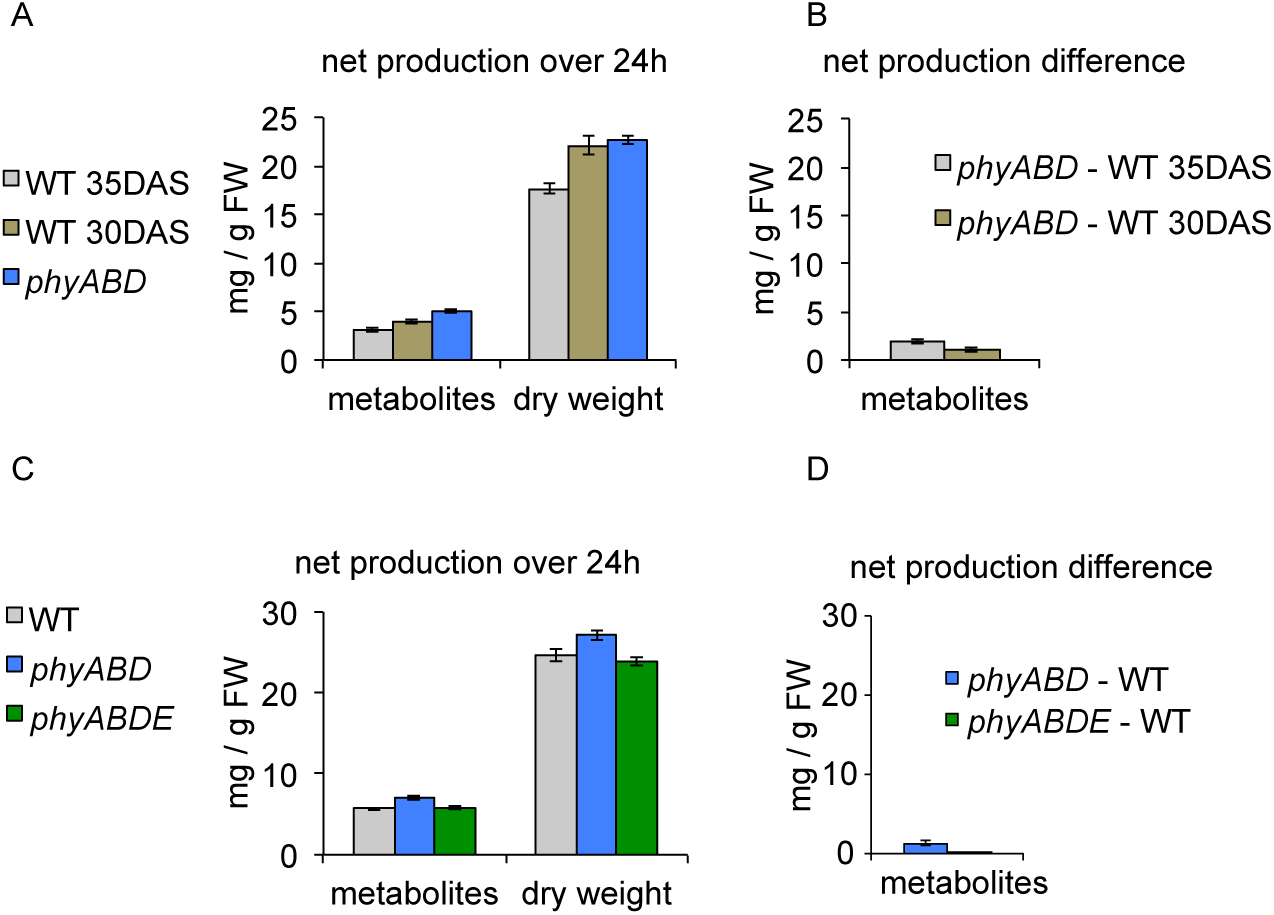
(A, C) Sum of key over-accumulated metabolites measured by enzymatic assay (glucose, fructose, sucrose, malate, fumarate, bulk amino acids, starch) gained (net gain) from EOD to EOD, calculated from relative growth rate and metabolite measurements per FW at ZT12. This was done for labelling experiment 1 (A) and 2 (C). (B, D) Difference in over-accumulated metabolites in (A) or (C) in *phy* mutants minus WT control(s), i.e. the net production difference, for labelling experiments 1 (B) and 2 (D). (B and D) are plotted on the same scaled as (a and c) to illustrate the biomass contained in the excess metabolites compared to the daily gain in dry biomass. errorbars: propagated SEM.

**Supplementary figure 11:**
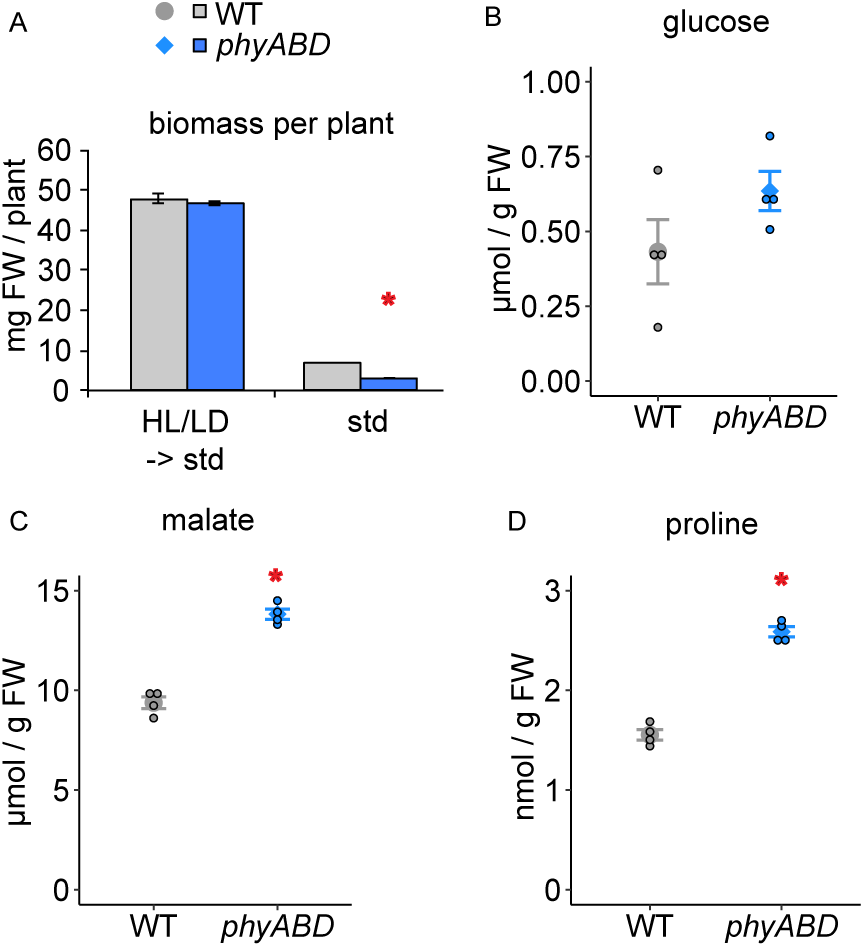
Biomass and metabolites in a condition that rescues the *phyABD* biomass deficit. WT and *phyABD* plants were grown in either our standard (‘std’) conditions (115µE, 12L:12D) for 18 DAS, or in higher light (HL) (350µE) and very long photoperiods (LD, 18L:6D) until 10 DAS, followed by 8 days of standard conditions (‘HL/LD -> std’). Biomass was measured in ‘std’ and ‘HL/LD -> std’ (A), as well as selected metabolites in ‘HL/LD -> std’ samples: (B) glucose, (C) malate, (D) proline. In (B-D) smaller round circles indicate values of individual replicates. Red asterisk: significant difference between *phyABD* and WT. Errorbars: SEM.

## References

Arrivault SA, Guenther M, Ivakov A, Feil R, Vosloh D, Van Dongen JT, Sulpice R, Stitt M 2009. “Use of Reverse-Phase Liquid Chromatography, Linked to Tandem Mass Spectrometry, to Profile the Calvin Cycle and Other Metabolic Intermediates in Arabidopsis Rosettes at Different Carbon Dioxide Concentrations.” Plant Journal 59 (5): 824–39

Casal, JJ. 2013. “Photoreceptor Signaling Networks in Plant Responses to Shade.” Annual Reviews of Plant Biology 64: 403–27.

Chen M, and Chory J. 2011. “Phytochrome Signaling Mechanisms and the Control of Plant Development.” Trends in Cell Biology 21 (11): 664–71.

Chew YH, Wenden B, Flis A, Mengin V, Taylor J, Davey CL, Tindal C, Thomas H, Ougham HJ, de Reffye P, Stitt M, Williams M, Muetzelfeldt R, Halliday KJ, and Millar AJ. 2014. “Multiscale Digital Arabidopsis Predicts Individual Organ and Whole-Organism Growth.” PNAS 112 (19): E3127–4136.

Cook D, Fowler S, Fiehn O, and Thomashow MF. 2004. “A Prominent Role for the CBF Cold Response Pathway in Configuring the Low-Temperature Metabolome of Arabidopsis.” PNAS 101 (42): 15243–48.

Devlin, PF, Robson PR, Patel SR, Goosey L, Sharrock RA, and Whitelam GC. 1999. “Phytochrome D Acts in the Shade-Avoidance Syndrome in Arabidopsis by Controlling Elongation Growth and Flowering Time.” Plant Physiology 119 (3): 909–15.

Figueroa CM, Feil R, Ishihara H, Watanabe M, Kölling K, Krause U, Höhne M, Encke B, Plaxton WC, Zeeman SC, Li Z, Schulze WX, Hoefgen R, Stitt M and Lunn JE. 2016. “Trehalose 6-Phosphate Coordinates Organic and Amino Acid Metabolism with Carbon Availability.” Plant Journal 85 (3): 410–23.

Figueroa, CM., and Lunn JE. 2016. “A Tale of Two Sugars: Trehalose 6-Phosphate and Sucrose.” Plant Physiology 172 (1): 7–27.

Flis A, Mengin V, Ivakov AA, Mugford ST, Hubberten HM, Encke B, Krohn N, Höhne M, Feil R, Hoefgen R, Lunn JE, Millar AJ, Smith AM, Sulpice R, Smith AM, Sulpice R, Stitt M. 2019. “Multiple Circadian Clock Outputs Regulate Diel Turnover of Carbon and Nitrogen Reserves.” Plant Cell and Environment 42 (2): 549–73.

Flis A, Sulpice R, Seaton D, Ivakov AA., Liput M, Abel C, Millar AJ, and Stitt M. 2016. “Photoperiod-Dependent Changes in the Phase of Core Clock Transcripts and Global Transcriptional Outputs at Dawn and Dusk in Arabidopsis.” Plant Cell and Environment 39 (9): 1955–81.

Fowler, S, and Thomashow MF. 2002. “Arabidopsis Transcriptome Profiling Indicates That Multiple Regulatory Pathways Are Activated during Cold Acclimation in Addition to the CBF Cold Response Pathway.” Plant Cell 14: 1675–90.

Franklin, KA., and Whitelam GC. 2005. “Phytochromes and Shade-Avoidance Responses in Plants.” Annals of Botany 96 (2): 169–75. 2007. “Light-Quality Regulation of Freezing Tolerance in Arabidopsis Thaliana.” Nature Genetics 39 (11): 1410–13.

Franklin KA, Praekelt U, Stoddart WM, Billingham OE, Halliday KJ, and Whitelam GC. 2003. “Phytochromes B, D, and E Act Redundantly to Control Multiple Physiological Responses in Arabidopsis.” Plant Physiology 131 (3): 1340–46.

Fukushima A, Kusano M, Nakamichi N, Kobayashi M, Hayashi N, Sakakibara H, Mizuno T, and Saito K. 2009. “Impact of Clock-Associated Arabidopsis Pseudo-Response Regulators in Metabolic Coordination.” PNAS 106 (21): 7251–56.

Gilmour SJ, Sebolt AM, Salazar MP, Everard JD, and Thomashow MF. 2000. “Overexpression of the Arabidopsis CBF3 Transcriptional Activator Mimics Multiple Biochemical Changes Associated with Cold Acclimation.” Plant Physiology 124 (4): 1854–65.

Gilmour SJ, Fowler SJ, and Thomashow MF. 2004. “Arabidopsis Transcriptional Activators CBF1, CBF2, and CHF3 Have Matching Functional Activities.” Plant Molecular Biology, 767–81.

Halliday KJ. 2003. “Changes in Photoperiod or Temperature Alter the Functional Relationships between Phytochromes and Reveal Roles for PhyD and PhyE.” Plant Physiology 131 (4): 1913–20.

Hare PD, and Cress WA. 1997. “Metabolic Implications of Stress-Induced Proline Accumulation in Plants.” Plant Growth Regulation 21: 79–102.

Hildebrandt TM. 2018. “Synthesis versus Degradation: Directions of Amino Acid Metabolism during Arabidopsis Abiotic Stress Response.” Plant Molecular Biology 98 (1–2): 121–35.

Hildebrandt, TM., Adriano Nunes Nesi, Wagner L. Araújo, and Hans Peter Braun. 2015. “Amino Acid Catabolism in Plants.” Molecular Plant 8 (11): 1563–79.

Hu W, Franklin KA, Sharrock RA, Jones M, Harmer SL, and Lagarias JC. 2013. “Unanticipated Regulatory Roles for Arabidopsis Phytochromes Revealed by Null Mutant Analysis.” PNAS 110 (4): 1542–47.

Huege J, Götze JP, Dethloff F, Junker B, and Kopka J. 2014. “Quantification of Stable Isotope Label in Metabolites via Mass Spectrometry.” Methods in Molecular Biology 1056: 213–23.

Hwang G, Kim S, Cho J-Y, Paik I, Kim J-I, and Oh E. 2019. “Trehalose-6-phosphate Signaling Regulates Thermoresponsive Hypocotyl Growth in Arabidopsis Thaliana.” EMBO Reports, 1–11. https://doi.org/10.15252/embr.201947828.

Ishihara H, Obata T, Sulpice R, Fernie AR, and Stitt M. 2015. Quantifying Protein Synthesis and Degradation in Arabidopsis by Dynamic 13CO2 Labeling and Analysis of Enrichment in Individual Amino Acids in Their Free Pools and in Protein. Plant Physiology. 168:74–93.

Johansson H, Jones HJ, Foreman J, Hemsted JR, Steward K, Grima R, and Halliday KJ. 2014. “Arabidopsis Cell Expansion Is Controlled by a Photothermal Switch.” Nature Communications 5:4848.

Kaplan F, Kopka J, Haskell DW, Zhao W, Schiller KC, Gatzke N, Sung DY, and Guy CL. 2004. “Exploring the Temperature-Stress Metabolome.” Plant Physiology 136: 4159–68.

Kempa S, Krasensky J, Santo SD, Kopka J, and Jonak C. 2008. “A Central Role of Abscisic Acid in Stress-Regulated Carbohydrate Metabolism.” PLoS ONE 3 (12): e3935.

Kim J, Kang H, Park J, Kim W, Yoo J, Lee N, Kim J, Yoon TY, and Choi G. 2016. “PIF1-Interacting Transcription Factors and Their Binding Sequence Elements Determine the in Vivo Targeting Sites of PIF1.” Plant Cell 28 (6): 1388–1405.

Kircher S, and Schopfer P. 2012. “Photosynthetic Sucrose Acts as Cotyledon-Derived Long-Distance Signal to Control Root Growth during Early Seedling Development in Arabidopsis.” PNAS 109 (28): 11217–21.

Kölling K, George GM, Künzli R, Flütsch P, and Zeeman SZ. 2015. “A Whole - Plant Chamber System for Parallel Gas Exchange Measurements of Arabidopsis and Other Herbaceous Species.” Plant Methods 11 (48).

Leivar P, Monte E, Oka Y, Liu T, Carle C, Castillon A, Huq E, and Quail PH. 2008. “Multiple Phytochrome-Interacting BHLH Transcription Factors Repress Premature Seedling Photomorphogenesis in Darkness.” Current Biology 18 (23): 1815–23.

Leivar P, Tepperman JM, Cohn MM, Monte E, Al-Sady B, Erickson E, and Quail PH. 2012. “Dynamic Antagonism between Phytochromes and PIF Family Basic Helix-Loop-Helix Factors Induces Selective Reciprocal Responses to Light and Shade in a Rapidly Responsive Transcriptional Network in Arabidopsis.” The Plant Cell 24 (4): 1398–1419.

Lilley-Steward JL, Gee CW, Sairanen I, Ljung K, and Nemhauser JL. 2012. “An Endogenous Carbon-Sensing Pathway Triggers Increased Auxin Flux and Hypocotyl Elongation.” Plant Physiology 160 (4): 2261–70.

Lisec J, Schauer N, Kopka J, Willmitzer L, and Fernie AR. 2006. “Gas Chromatography Mass Spectrometry-Based Metabolite Profiling in Plants.” Nature Protocols 1 (1): 387–96.

Lorrain S, Allen T, Duek PD, Whitelam GC, and Fankhauser C. 2008. “Phytochrome-Mediated Inhibition of Shade Avoidance Involves Degradation of Growth-Promoting BHLH Transcription Factors.” Plant Journal 53 (2): 312–23.

Lunn JE, Feil R, Hendriks JHM, Gibon Y, Morcuende R, Osuna D, Scheible W-R, Carillo P, Hajirezaei M-R, and Stitt M. 2006. “Sugar-Induced Increases in Trehalose 6-Phosphate Are Correlated with Redox Activation of ADPglucose Pyrophosphorylase and Higher Rates of Starch Synthesis in Arabidopsis Thaliana.” The Biochemical Journal 397 (1): 139–48.

Lunn, JE, Delorge I, Figueroa CM, Van Dijck P, and Stitt M. 2014. “Trehalose Metabolism in Plants.” Plant Journal 79 (4): 544–67.

Martìn G, Rovira A, Veciana N, Soy J, Toledo-Ortiz G, Gommers CMM, Boix M, Henriques R, Minguet EG, Alabadí D, Halliday KJ, Leivar P, and MonteE. 2018. “Circadian Waves of Transcriptional Repression Shape PIF-Regulated Photoperiod-Responsive Growth in Arabidopsis.” Current Biology 28: 311–18.

Maruyama K, Takeda M, Kidokoro S, Yamada K, Sakuma Y, Urano K, Fujita M, Yoshiwara K, Matsukura S, Morishita Y, Sasaki R, Suzuki H, Saito K, Shibata D, Shinozaki K, and Yamaguchi-Shinozaki K. 2009. “Metabolic Pathways Involved in Cold Acclimation Identified by Integrated Analysis of Metabolites and Transcripts Regulated by DREB1A and DREB2A.” Plant Physiology 150 (4): 1972–80.

Maruyama K, Sakuma Y, Kasuga M, Ito Y, Seki M, Goda H, Shimada Y, Yoshida S, Shinozaki K, and Yamaguchi-Shinozaki K. 2004. “Identification of Cold-Inducible Downstream Genes of the Arabidopsis DREB1A/CBF3 Transcriptional Factor Using Two Microarray Systems.” Plant Journal 38 (6): 982–93.

Mengin V, Pyl ET, Moraes TA, Sulpice R, Krohn N, Encke B, and Stitt M, 2017. “Photosynthate Partitioning to Starch in Arabidopsis Thaliana Is Insensitive to Light Intensity but Sensitive to Photoperiod Due to a Restriction on Growth in the Light in Short Photoperiods.” Plant, Cell & Environment. https://doi.org/10.1111/pce.13000.

Mizuno T, Oka H, Yoshimura F, Ishida K, and Yamashino T. 2015. “Insight into the Mechanism of End-of-Day Far-Red Light (EODFR)-Induced Shade Avoidance Responses in Arabidopsis Thaliana.” Bioscience, Biotechnology and Biochemistry 79 (12): 1987–94.

Oyama T, Shimura Y, and Okada K. 1997. “The Arabidopsis HY5 Gene Encodes a BZIP Protein That Regulates Stimulus-Induced Development of Root and Hypocotyl.” Genes and Development 11 (22): 2983–95.

Pagter M, Alpers J, Erban A, Kopka J, Zuther E, and Hincha DK. 2017. “Rapid Transcriptional and Metabolic Regulation of the Deacclimation Process in Cold Acclimated Arabidopsis Thaliana.” BMC Genomics 18 (731): 1–17.

Rausenberger J, Hussong A, Kircher S, Kirchenbauer D, Timmer J, Nagy F, Schäfer E, and Fleck C. 2010. “An Integrative Model for Phytochrome B Mediated Photomorphogenesis: From Protein Dynamics to Physiology.” PLoS ONE 5 (5): e10721.

Reed JW, Nagpal P, Poole DS, Furuya M, and Chory J. 2007. “Mutations in the Gene for the Red/Far-Red Light Receptor Phytochrome B Alter Cell Elongation and Physiological Responses throughout Arabidopsis Development.” The Plant Cell 5 (2): 147.

Sánchez-Lamas M, Lorenzo CD, and Cerdán PD. 2016. “Bottom-up Assembly of the Phytochrome Network.” PLoS Genetics 12 (11): 1–24.

Sengupta S, Mukherjee S, Basak P, and Majumder AL. 2015. “Significance of Galactinol and Raffinose Family Oligosaccharide Synthesis in Plants.” Frontiers in Plant Science 6:656.

Stewart JL, Maloof JL, and Nemhauser JL. 2011. “PIF Genes Mediate the Effect of Sucrose on Seedling Growth Dynamics.” PLoS ONE 6 (5): e19894.

Strasser B, Sánchez-Lamas M, Yanovsky MJ, Casal JJ, and Cerdán PD. 2010. “Arabidopsis Thaliana Life without Phytochromes.” PNAS 107 (10): 4776–81.

Sweetlove LJ, Beard KFM, Nunes-Nesi A, Fernie AR, and Ratcliffe RG. 2010. “Not Just a Circle: Flux Modes in the Plant TCA Cycle.” Trends in Plant Science 15 (8): 462–70.

Szecowka M, Heise R, Tohge T, Nunes-Nesi A, Vosloh D, Huege J, Feil R, Lunn JE, Nikoloski Z, Stitt M, Fernie AR, and Arrivault S. 2013. “Metabolic Fluxes in an Illuminated Arabidopsis Rosette.” The Plant Cell 25 (2): 694–714.

Taji T, Ohsumi C, Iuchi S, Seki M, Kasuga M, Kobayashi M, Yamaguchi-Shinozaki K, and Shinozaki K. 2002. “Important Roles of Drought- and Cold-Inducible Genes for Galactinol Synthase in Stress Tolerance in Arabidopsis Thaliana.” The Plant Journal 29 (4): 417–26.

Urano K, Maruyama K, Ogata Y, Morishita Y, Takeda M, and Sakurai N. 2009. “Characterization of the ABA-Regulated Global Responses to Dehydration in Arabidopsis by Metabolomics.” The Plant Journal 57: 1065–78.

Wang X, Wang L, Wang Y, Liu H, Hu D, Zhang N, Zhang S, Cao H, Cao Q, Zhang Z, Tang S, Song D, and Wang C. 2018. “Arabidopsis PCaP2 Plays an Important Role in Chilling Tolerance and ABA Response by Activating CBF- and SnRK2-Mediated Transcriptional Regulatory Network.” Frontiers in Plant Science 9:215.

de Wit M, George GM, Çaka Ince Y, Dankwa-Egli B, Hersch M, Zeeman SC, and Fankhauser C. 2018. “Changes in Resource Partitioning between and within Organs Support Growth Adjustment to Neighbor Proximity in Brassicaceae Seedlings.” PNAS 115 (42): E9953–61.

de Wit M, Ljung K, and Fankhauser C. 2015. “Contrasting Growth Responses in Lamina and Petiole during Neighbor Detection Depend on Differential Auxin Responsiveness Rather than Different Auxin Levels.” New Phytologist 208 (1): 198–209.

Xin Z, and Browse J. 1998. “Eskimo1 Mutants of Arabidopsis Are Constitutively Freezing-Tolerant.” PNAS 95: 7799–7804.

Yadav UP, Ivakov A, Feil R, Duan GY, Walther D, Giavalisco P, Piques M, Carillo P, Hubberten H-M, Stitt M and Lunn JE. 2014. “The Sucrose-Trehalose 6-Phosphate (Tre6P) Nexus: Specificity and Mechanisms of Sucrose Signalling By.” Journal of Experimental Botany 65 (4): 1051–68.

Yang D, Seaton DD, Krahmer J, and Halliday KJ. 2016. “Photoreceptor Effects on Plant Biomass, Resource Allocation, and Metabolic State.” PNAS 113 (27): 7667–72.

